# Depletion of the RNA-binding protein PURA triggers changes in posttranscriptional gene regulation and loss of P-bodies

**DOI:** 10.1101/2022.02.09.479353

**Authors:** Lena Molitor, Melina Klostermann, Sabrina Bacher, Juliane Merl-Pham, Nadine Spranger, Sandra Burczyk, Carolin Ketteler, Ejona Rusha, Daniel Tews, Anna Pertek, Marcel Proske, Anke Busch, Sarah Reschke, Regina Feederle, Stefanie M. Hauck, Helmut Blum, Micha Drukker, Pamela Fischer-Posovszky, Julian König, Kathi Zarnack, Dierk Niessing

**Author notes:** Correspondence should be addressed to: Kathi Zarnack, Dierk Niessing. Shared first authors (Lena Molitor, Melina Klostermann).

## Abstract

The RNA-binding protein PURA has been implicated in the rare, monogenetic, neurodevelopmental disorder PURA Syndrome. PURA binds both DNA and RNA and has been associated with various cellular functions. Only little is known about its main cellular roles and the molecular pathways affected upon *PURA* depletion. Here, we show that PURA is predominantly located in the cytoplasm, where it binds to thousands of mRNAs. Many of these transcripts change abundance in response to *PURA* depletion. The encoded proteins suggest a role for PURA in immune responses, mitochondrial function, autophagy and processing (P)-body activity. Intriguingly, reduced PURA levels decrease the expression of the integral P-body components LSM14A and DDX6 and strongly affect P-body formation in human cells. Furthermore, *PURA* knockdown results in stabilization of P-body-enriched transcripts, whereas other mRNAs decrease. Hence, reduced PURA levels, as reported in patients with PURA Syndrome, influence the formation and composition of this phase-separated RNA processing machinery. Our study proposes PURA Syndrome as a new model to study the tight connection between P-body-associated RNA regulation and neurodevelopmental disorders.

**Graphical Abstract:** 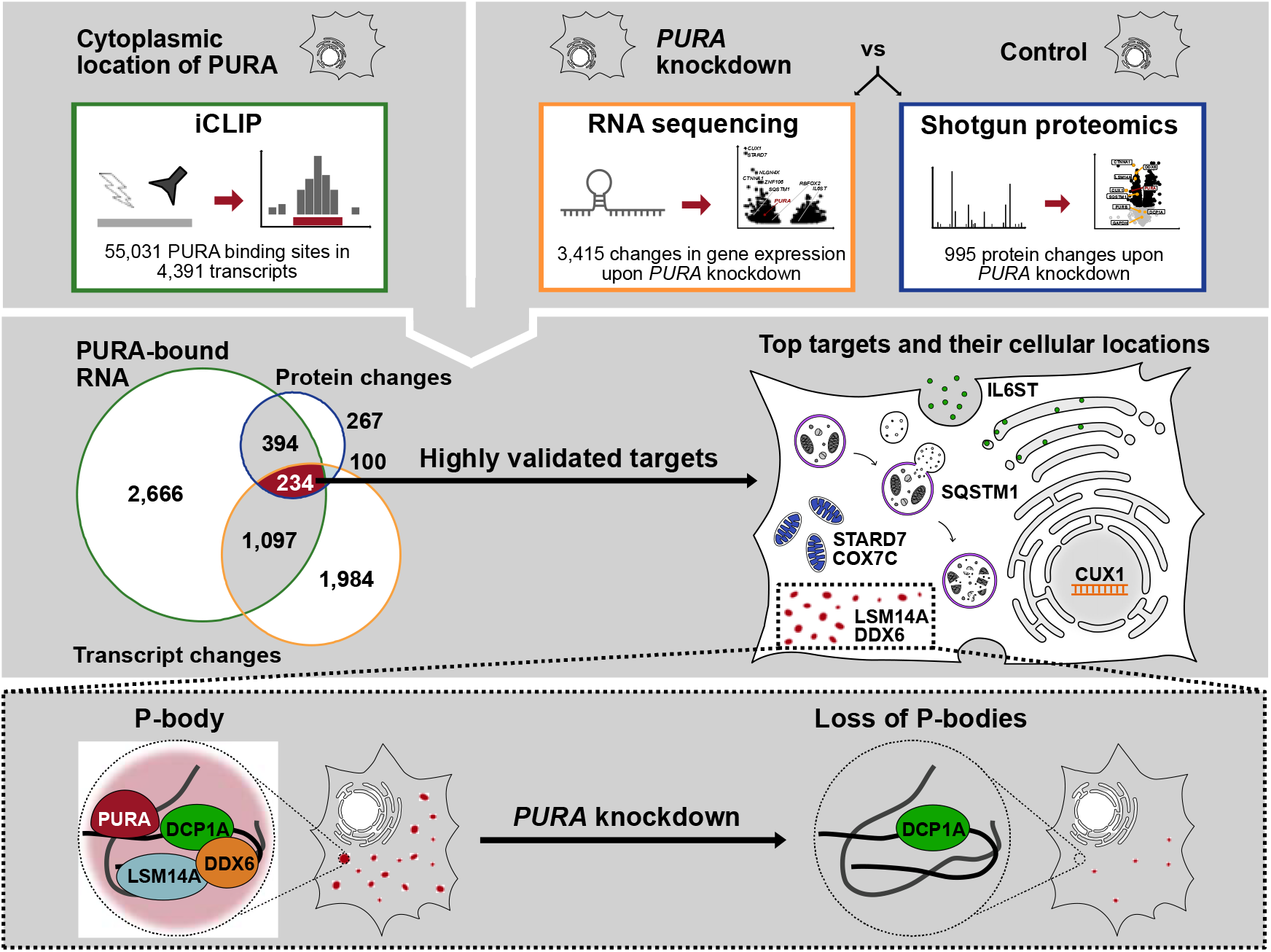

## Introduction

In recent years, an increasing number of RNA-binding proteins have been implicated in the pathology of human disorders, in particular in neuronal diseases (1,2). One of them is the purine rich element binding protein A (PURA, formerly known as Puralpha) (3). In 2014, mutations in the *PURA* gene have been linked to the neurodevelopmental disorder PURA Syndrome (4,5). Patients with heterozygous *de novo* mutations in *PURA* show a range of symptoms including neurodevelopmental delay, intellectual disability, hypotonia, and epilepsy (6,7). For unknown reasons, the phenotypes of patients with PURA Syndrome vary considerably. In addition, mice with homozygous deletions of *PURA* from two independent studies show phenotypes partially resembling patients with PURA Syndrome, including tremor and movement abnormalities (8,9). While it is generally assumed that symptoms are caused by haploinsufficiency of the PURA protein (10,11), the affected molecular pathways remain largely elusive. Furthermore, mice with a heterozygous mutation in *PURA* show very little overlap with human symptoms (12), underlining the importance of PURA-related studies in human cells.

PURA was initially reported as a transcription factor binding to a specific DNA sequence motif upstream of the *c-myc* gene (13,14). While this and other DNA targets of PURA have been described in the past (15), the majority of recent reports indicated an involvement in RNA-related processes. In particular, PURA was repeatedly found in cytoplasmic, kinesin- and myosin-containing mRNA-transport complexes in neurons of mice (16-20). Furthermore, PURA was reported to localize to cytoplasmic stress granules in human cells. In a cell culture model for amyotrophic lateral sclerosis (ALS), changes in PURA expression modulated the pathological appearances of stress granules and mitigated ALS-related neurotoxicity, indicating a functional importance of PURA for these membrane-less organelles (21). While these reports indicate that PURA associates with both DNA and RNA, it remains unclear which interaction is functionally more relevant.

PURA belongs to the PC4 family of proteins (22) that bind single-stranded (ss) nucleic acids. This family of proteins has been found in all kingdoms of life and often show similarities in their nucleic acid binding properties. Nuclear magnetic resonance (NMR) measurements and quantitative binding studies revealed that PURA interacts with ssDNA and ssRNA with comparable affinities and properties (23). Crystal structures of PURA proteins from different species revealed two so-called PUR domains, which mediate PURA’s dimerization, and indicated that there are no structural constraints favoring DNA or RNA binding (23,24). Thus, although we have a relatively clear understanding of PURA’s structural and molecular properties (3), a link is missing that explains how its nucleic acid binding impacts cellular functions. Hence, the aim of this study was to identify target pathways of PURA that could be dysregulated in patients with PURA Syndrome.

## Material and Methods

### Generation of a PURA-specific antibody

The monoclonal IgG2-2a anti-PURA^12D11^ antibody was raised against the unstructured linker region (21 amino acids) between PUR I-II of human PURA (peptide sequence: Cys-AQLGPSQPPDLAQAQDEPRRA; Peps4LS GmbH, Heidelberg) (**Supplementary Figure S1**). Several antibodies were generated after immunization of Sprague Dawley rats with ovalbumin-coupled peptide and hybridoma fusion as described before (25). Antibody supernatants were validated for binding to biotinylated PURA peptide in a solid-phase immunoassay. Subsequently, the panel of antibody supernatants was tested by Western blot on total cell lysate from HeLa cells. Selected clones were subcloned twice by limiting dilution to obtain stable monoclonal cell lines. Finally, the best antibody was tested in Western blot experiments on total cell lysate from *PURA* knockdown (KD) and control (CTRL) HeLa cells.

Cross-reactivity to PURB was excluded using recombinantly expressed GST-PURA and GST-PURB in Western blot assays (**Supplementary Figure S1G**). For this, human PURA and PURB were cloned into the pOPIN-J expression vector and expressed in *E. coli* Rosetta 2 (DE3) cells, followed by Western blot with cell lysates. Experiments in this work were performed with hybridoma supernatant clone 12D11 (IgG2a/⍰). For additional details, see “Western blot experiments”.

### Generation of inducible PURA overexpression HeLa cell line

In the overexpression construct GFP-P2A-FLAG-PURA, GFP and FLAG-PURA are separated by a viral P2A cleavage site which mediates translation into two disconnected protein products (26). PiggyBac plasmids (27) harboring GFP-P2A-FLAG-PURA under control of a doxycycline-inducible promoter were generated using Gibson assembly (GeneArt Thermo Fisher Scientific) of three fragments containing overhangs according to manufacturer’s instructions. Using these plasmids, overexpression cell lines were generated. The plasmid containing GFP-P2A-FLAG-PURA as well as a PiggyBac Helper plasmid (27) were transfected into cells using Lipofectamine 3000 (Thermo Fisher Scientific) according to manufacturer’s instructions. Transfected cells were incubated at 37 °C, 5% CO_2_ for two to four days. Transfected cells were subsequently selected by addition of hygromycin (700 μg/ml) for at least ten days. GFP-P2A-FLAG-PURA overexpression was induced by addition of doxycycline (1 μg/ml) for 24 hours (h). Successful translation of the overexpression construct was validating by showing GFP+ cells using GFP fluorescence, while the overexpressed FLAG-PURA protein was detected using the anti-PURA^12D11^ antibody.

### siRNA-mediated *PURA* knockdown

siRNA-mediated knockdown of *PURA* in HeLa cells was performed using a predesigned *PURA* siRNA pool (Dharmacon, M-012136-01-000). One day prior to transfection, HeLa cells were plated at 20% confluency in 6-well plates. The next day, lipofection was performed using RNAiMAX (Thermo Fisher Scientific) according to manufacturer’s instructions. Briefly, 20 μM siRNA pool was mixed with OptiMEM (Thermo Fisher Scientific) in one reaction and lipofectamine RNAiMAX was mixed with OptiMEM in a second reaction. Both reactions were incubated at room temperature (RT) for 5 min and mixed by flicking. Meanwhile, culture media on HeLa cells was changed to 1 ml DMEM + 10% FBS lacking antibiotics. Contents of both reactions were mixed and further incubated for 20 min at RT. Finally, 400 μl of lipofection mix were added drop-wise onto HeLa cells. As a control, siGENOME Non-Targeting Pool #1 (Dharmacon, D-001206-13-05) and a second independent predesigned *PURA* siRNA (Thermo Fisher Scientific, 289567) was used as described for the *PURA* siRNA pool. The transfected cells were incubated for 48 h at 37 °C and 5% CO_2_ and subsequently analyzed in downstream assays.

### RNA sequencing

Four independent biological replicates of *PURA* KD (siGENOME Non-Targeting Pool #1; Dharmacon, D-001206-13-05) and CTRL HeLa cells were prepared as described above. Control cells were only treated with lipofectamine RNAiMAX (Thermo Fisher Scientific) and OptiMEM (Thermo Fisher Scientific) without adding the non-targeting pool siRNA. After 48 h incubation with siRNA, cells were washed using PBS and subsequently collected in TRIzol Reagent (Thermo Fisher Scientific). Total RNA was extracted using Maxwell^®^ RSC miRNA Tissue Kit (AS1460) on the Maxwell^®^ RSC instrument according to the manufacturer’s instructions. Briefly, shock-frozen cells were resuspended in 500 μl Trizol. 6 μl 1-thioglycerol were added to 300 μl Trizol suspension. The suspension was mixed vigorously and total RNA extracted by phase separation using RSC SimplyRNA Tissue (AS1340) setup. The concentration was measured on a NanoDrop spectrometer (Thermo Fisher Scientific) and the quality of the RNA samples checked on an Agilent Bioanalyzer (Agilent; RNA 6,000 Nano kit).

Libraries were generated with the Lexogen SENSE mRNA-seq Library Prep Kit V2 (Lexogen) with half of the volume described in the manual. Briefly, 100 ng of total RNA were bound to oligo-dT beads, reverse-transcribed on the beads and the second strand of cDNA was synthesized after ligation of an adapter. The eluted cDNA was amplified with barcoded primers during 11 cycles of PCR with the following program: 98 °C for 10 s, 65 °C for 20 s, 72 °C for 30 s and a final extension at 72 °C for 1 min (hold at 10 °C). The PCR products were purified with Agencourt AMPure XP beads (Beckman Coulter). The quality of the libraries was validated with an Agilent Bioanalyzer (DNA 1,000 Kit). The finished, barcoded libraries were pooled and sequenced on an Illumina HiSeq1500 as 50-nt single-end reads. Each library yielded on average 35 million reads.

Reads were aligned to the human genome (GRCh38.p12 from GENCODE) with STAR (version 2.7.6a) (28) with up to 4% mismatches and no multimapping allowed. Reads per gene were counted using htseq-count (version 0.11.3) (29) with default settings and GENCODE gene annotation (release 31) (30). Differential expression analysis between *PURA* KD and CTRL samples was performed using DESeq2 (version 1.33.4) (31) with a significance cutoff at an adjusted *P* value < 0.01 (Benjamini-Hochberg correction). This yielded a total of 3,415 significantly differentially expressed transcripts, including 1,663 with increased and 1,752 with decreased expression upon *PURA* KD. Note that we use the term transcripts equivalent to genes here. For the heatmap in **Figure 4C**, library size-corrected RNA-seq read counts per sample were rlog-transformed (DESeq2, version 1.33.4) (31). For visualization, the counts were converted into row-wise *z*-scores.

### Shotgun proteomics

#### Generation of protein lysates

Four independent biological replicates of *PURA* KD and CTRL HeLa cells were prepared as described above. Cell samples were collected by scraping them in PBS and subsequent centrifugation at 500 x g for 3 min. Cell pellets were lysed in 100 μl RIPA buffer and subsequently centrifuged at 16,000 x g at 4 °C for 10 min. The supernatant was transferred to a new tube and the amount of protein in the lysate was determined by Bradford assay (ROTI^®^Quant, Carl Roth).

#### Sample preparation for mass spectrometric analysis

10 μg per sample were digested with Lys-C and trypsin using a modified FASP procedure (32,33). Briefly, after reduction and alkylation using DTT and IAA, the proteins were centrifuged on a 30 kDa cutoff filter device (Sartorius) and washed each thrice with UA buffer (8 M urea in 0.1 M Tris/HCl pH 8.5) and with 50 mM ammonium bicarbonate. The proteins were digested for 2 h at RT using 0.5 μg Lys-C (Wako Chemicals) and for 16 h at 37 °C using 1 μg trypsin (Promega). After centrifugation (10 min at 14 000 g), the eluted peptides were acidified with 0.5% TFA and stored at -20 °C.

#### LC-MS/MS measurements

Liquid chromatography with tandem mass spectrometry (LC-MS/MS) analysis was performed on a QExactive HFX mass spectrometer (Thermo Fisher Scientific) online coupled to a UItimate 3000 RSLC nano-HPLC (Dionex). Samples were automatically injected and loaded onto the C18 trap cartridge and after 5 min eluted and separated on the C18 analytical column (Acquity UPLC M-Class HSS T3 Column, 1.8 μm, 75 μm x 250 mm; Waters) by a 90 min non-linear acetonitrile gradient at a flow rate of 250 nl/min. MS spectra were recorded at a resolution of 60,000 with an AGC target of 3 × 10e6 and a maximum injection time of 30 ms from 300 to 1500 m/z. From the MS scan, the 15 most abundant peptide ions were selected for fragmentation via HCD with a normalized collision energy of 28, an isolation window of 1.6 m/z, and a dynamic exclusion of 30 s. MS/MS spectra were recorded at a resolution of 15,000 with a AGC target of 10e5 and a maximum injection time of 50 ms. Unassigned charges, and charges of +1 and >8 were excluded from precursor selection.

#### Quantitative MS analysis

Acquired raw data was analyzed in the MaxQuant software (MPI Biochemistry, Martinsried; version 1.6.7.0, (34) for peptide and protein identification via a database search (Andromeda search engine, (35) against the SwissProt Human database (Release 2020_02, 20435 sequences; 11,490,581 residues), considering full tryptic specificity, allowing for up to one missed tryptic cleavage sites, precursor mass tolerance 10 ppm, fragment mass tolerance 0.02 Da. Carbamidomethylation of cysteine was set as a static modification. Dynamic modifications included deamidation of asparagine and glutamine and oxidation of methionine. Identifications were filtered for a PSM false discovery rate < 1% and protein false discovery rate of 5%. Label-free quantifications were based on unique peptides applying the LFQ algorithm (36) with LFQ min, count of 1 in combination with the default match-between runs settings, allowing for matching of identifications (37) between the individual runs resulting in quantifications of MS features throughout the dataset.

Differential protein expression was analyzed based on MaxQuant LFQ values using DEqMS (1.12.1) (38). We treated missing values as follows: If protein abundance was not measured in any of the samples in one condition it was set to zero, which results in infinite values for the calculated ratios. Peptides with two or more missing values in one condition were treated as non-quantifiable and no ratios were calculated. When only one value was missing per condition, the ratios were calculated on the other three samples.

*P* values were adjusted for multiple testing by Benjamini-Hochberg correction. An advantage of DEqMS is that it implements scaling of (adjusted) *P* values to the number of detected unique peptides. We deemed proteins with a scaled adjusted *P* value < 0.05 significant. In the heatmap in **Figure 4C**, protein abundance is shown after z-score normalization.

### Quantitative real-time PCR (qPCR)

RNA was extracted using the High Pure RNA isolation kit (Roche Molecular Systems) according to manufacturer’s instructions and following the general precautions required for RNA work (39). RNA was eluted in 50 μl DEPC water. Two rounds of DNase digestion using TURBO DNA-free kit (Thermo Fisher Scientific) according to manufacturer’s instructions were performed to safely remove all remaining DNA. The amount of pure RNA was analyzed by measuring the OD_260_ using a NanoDrop microliter photometer. The extracted RNA was reverse transcribed using PrimeScript RT Master Mix (Takara) according to manufacturer’s instructions. On the generated cDNA libraries, qPCR experiments were performed using SYBR Green Master Mix (Thermo Fisher Scientific) in a LightCycler 480 Instrument II (Roche Molecular Systems) in 96-well format. c_t_ values were used for the analysis of differential mRNA abundance between *PURA* KD and CTRL conditions. This analysis was done using the ΔΔc_t_ method (40) with the housekeeping genes *GAPDH* and *RPL32* for normalization. Changes were tested for significance using an unpaired two-sided Student’s *t*-test on the ΔΔc_t_ values. All oligonucleotides used for qPCR experiments are listed in **Supplementary Table S8**.

### Western blot experiments

Western blot experiments were performed using Mini Blot module (Thermo Fisher Scientific) according to manufacturer’s instructions. NuPage loading dye (Thermo Fisher Scientific) was added onto cell lysates and the mix was heated to 70 °C for 10 min prior to gel loading. The blotting module was filled with NuPAGE 1x SDS running buffer (Thermo Fisher Scientific) and 500 μl NuPAGE antioxidant (Thermo Fisher Scientific) was added to the front part of the chamber. Samples were cooled to RT and loaded onto NuPAGE 4-12% Bis-Tris pre-cast gradient gels (Thermo Fisher Scientific) in the buffer-filled blotting module. For protein molecular weight estimation, BlueStar Plus Prestained Protein Marker (Nippon Genetics) was used. The gel was run for 45 min at 200 V. The PVDF blotting membrane was activated in methanol for 30 s prior to usage. All other components were soaked in 1x transfer buffer (Thermo Fisher Scientific) with 20% methanol prior to blotting module assembly. The assembled module was inserted into the Mini Blot module (Thermo Fisher Scientific), the surrounding chamber was filled with MQ water and the blotting reaction was run for 90 min at 30 V. Subsequently, the membrane was blocked using 1% casein in PBST. This was incubated for 30 min at RT while rotating on tube revolver machine (neoLab). Then, the blocking solution was discarded and primary antibody (dilutions in **Supplementary Table S11**) in 0.5% casein in PBST was added. Primary antibody was incubated overnight at 4 °C rotating on a tube revolver device. On the next day, the primary antibody solution was discarded, and blot was washed. Next, the secondary antibody (dilutions in **Supplementary Table S12**) in PBST was incubated on the membrane for 1 h at RT while rotating. Finally, the secondary antibody solution was discarded, and the membrane was washed. Then, the membrane was incubated with Amersham ECL Prime Western Blotting Detection Reagent (GE Healthcare). 1:1 mix of solution A and solution B of the above-mentioned ECL kit was added onto the membrane. The membrane was analyzed using Fusion SL 4 device (Vilber Lourmat, software FusionCapt Advance SL 4 16.04). Changes in signal intensity between *PURA* KD and CTRL samples were tested for significance in R using a paired one-sided Student’s *t*-test. For visualization, all samples were normalized to the mean of CTRL samples.

### Immunofluorescence staining

Cells were cultivated in culture medium on coverslips (Thermo Fisher Scientific) in 12-well plates and subsequently washed using PBS and fixed using 3.7% formaldehyde in PBS. Fixing media was incubated for 10 min at RT in the cell culture dish. Permeabilization was induced by incubating the cells in 0.5% Triton X-100 in PBS for 5 min at RT. Afterwards, cells were washed twice using PBS and the coverslips were transferred to a pre-labelled parafilm that was placed onto wet Whatman paper to prevent drying. Cells were then blocked for 10 min in blocking buffer (1% donkey serum in PBST) and incubated with primary antibodies (**Supplementary Table S11**) in blocking buffer for 1 h at RT. Secondary antibodies (**Supplementary Table S12**) were diluted in blocking buffer and incubated for 1 h at RT. Washing steps after antibody incubation were performed with PBST. DNA was stained with DAPI at 0.5 μg/ml in PBS by incubation for 5 min at RT, followed by washing twice using PBS. Finally, cells were mounted in ProLong Diamond Antifade (Thermo Fisher Scientific) and imaging was performed by fluorescence microscopy using an Axio.Observer.Z1 (Carl Zeiss AG) microscope. Immunofluorescence staining for P-bodies using the marker proteins DCP1A and LSM14A was performed on a LSM 880 Airyscan confocal microscope (Carl Zeiss AG) using a Zeiss Plan-Apochromat 20x/0.8 M27 objective and ZEN Black software (version 14.0.18.201).

### Quantitative analysis of immunofluorescence images

Quantification of DCP1A, LSM14A and DDX6 granules in different cell lines (HeLa and NHDF) was done using Fiji software (Version 2.3.0/1.53q). For each image, the cell number was determined using the auto threshold function on the DAPI channel, followed by particle analysis selecting a particle size of 1000-infinity pixel units (μm^2^). Then, all the granules on the image were quantified using the same particle analysis tool, after processing the corresponding channel with auto threshold. Granules were counted by selecting a particle size of 40-200 pixel units (μm^2^) and a circularity of 0.1-1.00 for HeLa cells (**Figure 5B-D**) (41,42). In NHDF cells, a particle size of 30-100 pixel units (μm^2^) was selected (**Supplementary Figure S10G, H**). Quantification was done in three biological replicates and for each replicate, four confocal pictures (20x magnification) of each condition (CTRL, *PURA* KD) were analyzed.

For the analysis of signal intensity of IL6ST and PURA immunofluorescence images, the channel for quantification was split from the DAPI channel. For each image, the background noise was removed using the subtract background tool with a rolling ball radius of 100 pixels. The rectangle tool was used to select parts of the picture without any signal and then that area was measured for integrated intensity. Afterwards the whole picture was measured with the same tool. For analysis, the background integrated intensity was subtracted from the integrated intensity of the channel of interest. The intensity of each image was also normalized to the cell number using the same particle analysis tool as described above (**Figure 4J, K, Supplementary Figure S9G**).

Colocalization was determined using a plugin from the Fiji software named BioIP JaCoP (43). The two channels to be analyzed were merged and then the plugin was used with a set threshold for Channel A and B “Otsu” (**Figure 5D**).

### Nuclear-cytoplasmic fractionation

5 million HeLa cells were utilized for cell fractioning. Cytoplasmic and nuclear fractions were generated using the NE-PER Kit (Thermo Scientific, Cat.-No: 78833) following the manufacturer’s instructions. In short, HeLa cells were resuspended, vortexed and centrifuged in cytoplasmic extraction buffer to obtain the cytoplasmic fraction. Afterwards, the pellet consisting of nuclei was resuspended in nuclear extraction buffer and frequently vortexed every 10 minutes for a total of 40 minutes to degrade the nuclear membrane. After centrifugation, the total protein concentration of the cytoplasmic and nuclear fraction were determined using BCA assay (Pierce™ BCA Protein Assay Kit, Cat.-No. 23227). For subcellular localization analysis of PURA, 20 μg of total protein for each cell fraction were used for western blotting. To validate successful separation of cell fractions protein levels of PARP1 (nuclear control: Sigma-Aldrich, Cat.-No. HPA045168) and GAPDH (cytoplasmic control: Sigma-Aldrich, Cat.-No. HPA040067) were additionally analysed **(Figure 1B, C)**.

**Figure 1.**
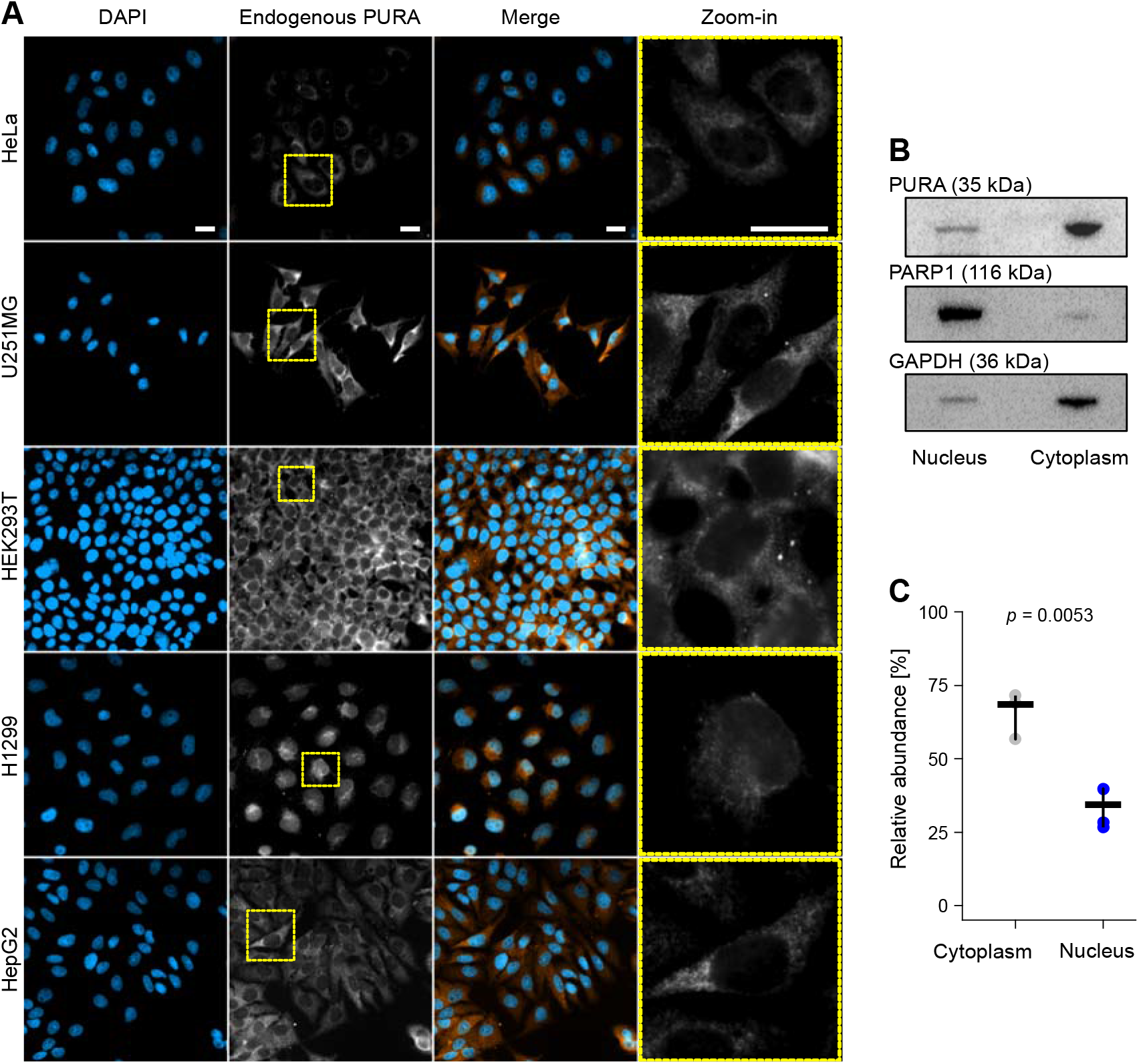
PURA is predominantly located in the cytoplasm. **(A)** Immortalized cell lines from different tissue origins were analyzed for PURA’s subcellular localization by immunofluorescence (IF) staining. Cells were stained with DAPI (blue) and anti-PURA^12D11^ (white). In the merged image, PURA staining is shown in orange for better visualization. Scale bars, 20 μm. **(B)** Nuclear-cytoplasmic fractionation of HeLa cell lysates. Representative Western blot using anti-PARP1 antibody (116 kDa) as marker for nuclear fraction, anti-GAPDH antibody as marker for cytoplasmic fraction (36 kDa) and anti-PURA^12D11^. **(C)** Densitometric quantification of PURA levels in cell fractions in 3 replicates (of which one is shown in (B)). Bars represent the mean ± standard deviation (SD) of three independent experiments, *P* value is calculated by unpaired two-sided Students *t*-test.

### Electrophoretic mobility shift assay (EMSA)

For EMSAs, GST-PURA (pOPIN-J) was recombinantly expressed in *E. coli* Rosetta 2 (DE3). Proteins were purified via GSTrap FF column (Cytiva). The GST-Tag was removed via 3C protease cleavage and the sample passed through HiTrap Q FF and HiTrap Heparin FF column. Protein bound to the HiTrap Heparin column was eluted and the sample further purified using HiLoad 16/600 Superdex 75 size-exclusion chromatography column (Cytiva) (**Supplementary Figure S3A-C**).

RNAs for EMSAs were *in vitro* transcribed using the MEGAshortscript T7 Transcription kit (Ambion). As a template, purchased HPLC-purified primers were used (**Supplementary Table S10**). T7 primer and the DNA template were annealed by incubation at 60 °C for 5 min and cooled down to RT. The *in vitro* transcription was performed as described by manufacturer (Ambion). Afterwards, the remaining DNA was digested using DNase I (Ambion, same kit) and the RNA was purified by phenol/chloroform extraction. Ethanol precipitation was performed at -20 °C with the addition of GlycoBlueTM coprecipitant (Thermo Fisher Scientific) and centrifugation for 1 min at 4 °C. DEPC water was used to dissolve the RNA and its quality was confirmed by denaturing and native polyacrylamide gel electrophoresis (PAGE).

For labeling of *in vitro* transcribed RNA, RNase-free buffers, materials, and reagents were used. *In vitro* transcribed RNA (15 pmol) was 5’ dephosphorylated using FastAP thermosensitive alkaline phosphatase (Thermo Fisher Scientific) in 20 μl reaction containing 1x FastAP buffer (Thermo Fisher Scientific) and 20 U of the RNase inhibitor SUPERaseIn (Thermo Fisher Scientific). The reaction was incubated for 30 min at 37 °C and the dephosphorylated RNA was extracted using phenol/chloroform and precipitated with 1 V 3 M NaOAc, 3 V absolute ethanol and cooled down at -20 °C for at least 15 min. For radioactive labeling, 15 pmol dephosphorylated RNA was 5’ phosphorylated with 32P from γ-32P ATP (Hartmann Analytic) with T4 PNK (New England Biolabs) in 1x PNK buffer in a 20 μl reaction. The reaction was incubated for 30 min at 37 °C and afterwards stopped for 10 min at 72 °C. NucAwayTM Spin column kit (Ambion) was used to remove unlabeled free nucleotides based on the manufacturer’s instructions. Eluted radiolabeled RNA was diluted to a final concentration of 100 nM in DEPC water and stored at -20 °C.

EMSAs for PURA-RNA interactions were performed as described before (23). The protein-nucleic acid complexes were formed in RNase-free binding buffer containing 250 mM NaCl, 20 mM Hepes pH 8.0, 3 mM MgCl_2_, 4 % glycerol, and 2 mM DTT. Serial protein dilutions and a constant amount of radiolabeled RNA (2.5 nM) were incubated in a total reaction volume of 20 μl for 20 min at RT. All experiments contained 100 μg/ml yeast tRNA (Ambion) as competitor. 10 μl of the reactions were loaded onto 6 % TBE polyacrylamide gels that were prepared beforehand. After electrophoresis (45 min, 100 V), gels were incubated for 15 min in fixing solution ([v/v] 10% acetic acid, [v/v] 30% methanol), dried in a gel dryer (BioRad) and analyzed with radiograph films using a Phosphoimager (Fujifilm VLA-3000) **(Figure 2D, Supplementary Figure S3D-M)**.

**Figure 2.**
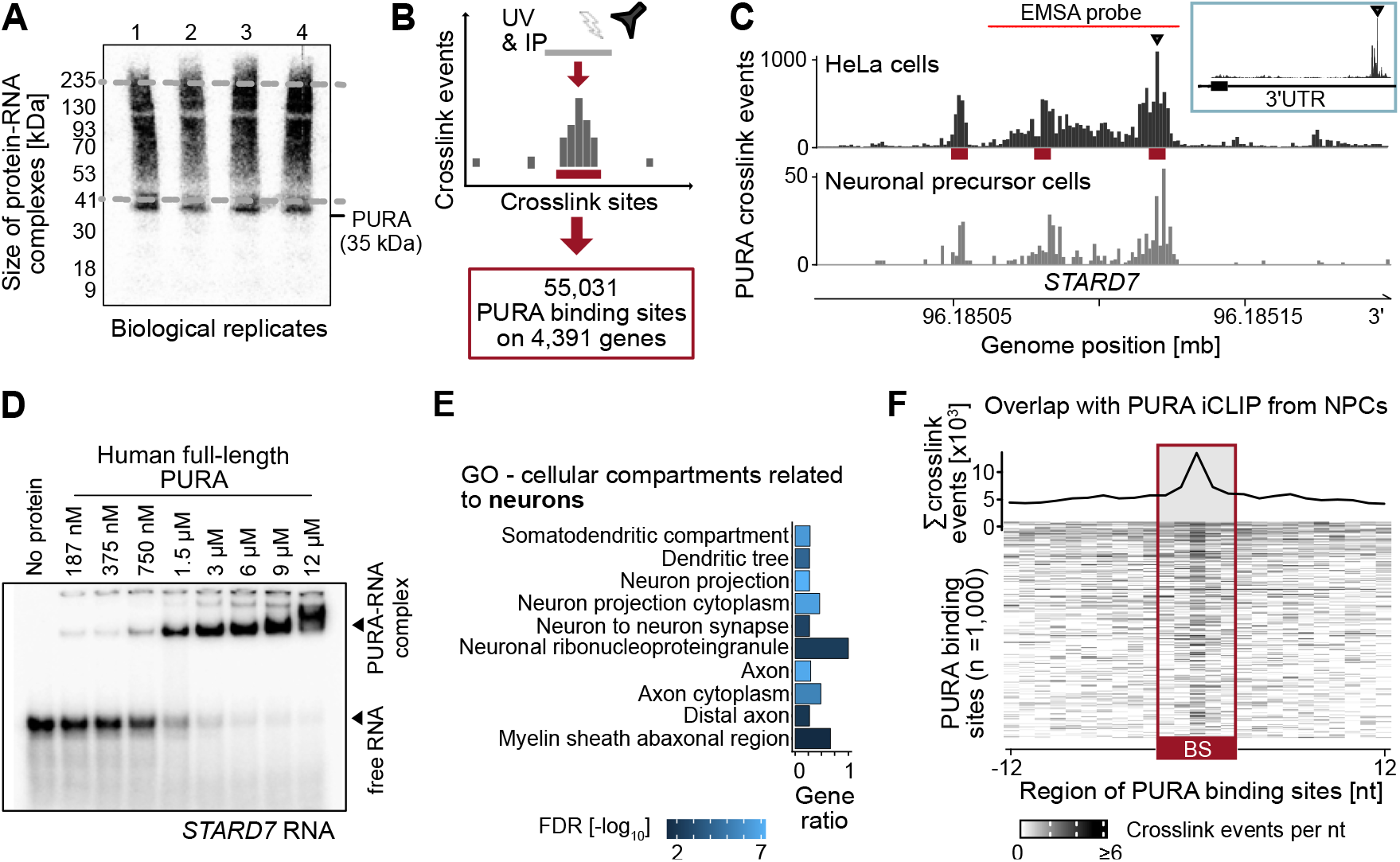
PURA globally binds RNAs via specific binding sites and regulates a subset of its target transcripts. **(A)** Autoradiogram with radiolabeled RNA after immunoprecipitation of endogenous PURA (anti-PURA^12D11^) from HeLa cells shows crosslinked PURA-RNA complexes. Co-purified RNA fragments of variable lengths result in a smear upstream of the expected molecular weight of PURA (35 kDa). Four biological replicates are shown. **(B)** Schematic depiction of iCLIP experiment that identified 55,031 binding sites for endogenous PURA in HeLa cells. **(C)** Genome browser view shows the pileup of crosslink events of endogenous PURA from HeLa cells (top) and neural progenitor cells (bottom) at binding sites (red) in the 3’UTR of the *STARD7* transcript. Inlay (blue box) shows complete *STARD7* 3’UTR. Red line denotes RNA fragment used for EMSA validation in (D). **(D)** EMSA with recombinantly expressed human full-length PURA and increasing concentrations (187 nM to 12 μM) of a radiolabeled 54-nt RNA fragment harboring two PURA binding sites from the *STARD7* 3’UTR (marked in (C)) confirms direct binding *in vitro*. For more EMSA experiments, see **Supplementary Figure S3. (E)** Cellular compartments (Gene Ontology) enriched in PURA-bound transcripts that relate to neurons (false discovery rate [FDR] < 0.05). Gene ratio (x-axis) depicts ratio of PURA-bound transcripts over all expressed transcripts (transcripts per million [TPM] > 0) attributed to this term. FDR is displayed as log_10_ by color scale. For a complete list of significantly enriched terms, see **Supplementary Table S3. (F)** PURA crosslink events from neural progenitor cells (NPCs) enrich in PURA binding sites identified in HeLa cells. Metaprofile (top) and heatmap (bottom) display crosslink events per nucleotide of endogenous PURA in NPCs for 1,000 binding sites with highest signal in the NPC iCLIP data. BS, binding site.

### Neural progenitor cell derivation from human induced pluripotent stem cells

The protocol for differentiation of neural progenitor cells was based on the generation of neurospheres (44). Briefly, the human induced pluripotent stem cell (hiPSC) line HMGU12 was harvested using StemMACS Passaging Solution (Miltenyi Biotec) and resuspended in DMEM/F-12 medium supplemented with 20% KSR, 1% NEAA, 1% GlutaMAX (Thermo Fisher Scientific), 10 μM SB431542, 5 μM dorsomorphin, 3 μM CHIR99021, 10 μM purmorphamine (Miltenyi Biotec), and 10 μM Y-27632 (Bio-Techne). The suspension was plated on an ultra-low attachment 6-well plate (Corning). Fresh medium was applied 24 h later without Y-27632. 48 h later, the basal medium was exchanged with N2B27-based medium containing a 1:1 mixture of DMEM-F-12 and Neurobasal A supplemented with 0.5% N-2, 1% B-27 minus vitamin A, 1% NEAA, and 1% GlutaMAX (Thermo Fisher Scientific), and the above-described small molecules. On day 5, N2B27-based medium was supplemented only with 50 μg/ml L-ascorbic acid, SB431542, and dorsomorphin. Approximately 24 h prior to replating, the medium was supplemented additionally with 5 ng/ml bFGF (Peprotech). On day 8, the neurospheres were mechanically dissociated and plated 1:6 on Matrigel-coated plates. Plated neurospheres were maintained for 7 days on N2B27 basal medium supplemented with Dual Smad inhibitors (Miltenyi Biotec (SB), Tocris (DM)), ascorbic acid and bFGF. On day 14, confluent neuroepithelial outgrowths were passaged in a 1:10 dilution using collagenase IV. The resulting neural progenitor cell (NPC) cultures were passaged every 7 days and maintained in N2B27 basal medium as described above with medium change every other day.

### iCLIP experiments

iCLIP experiments for PURA were performed according to the published iCLIP2 protocol (45) with minor adaptions. Cells were grown to confluence, washed using PBS, crosslinked using Stratalinker 2400 (Vilber Lourmat) at 254 nm and 300 mJ/cm^2^ and lysed. Immunoprecipitation of PURA from cell lysates was either performed using anti-PURA^12D11^-coated Dynabeads protein G (Thermo Fisher Scientific) or FLAG M2 beads (Sigma Aldrich). Subsequently, co-purified RNAs were dephosphorylated at the 3’ end, the 3’ adapter was ligated and the 5’ end was labeled with radioactive isotopes (γ-^32^P ATP; Hartmann Analytic). Samples were loaded onto a NuPAGE 4-12% Bis-Tris pre-cast SDS gradient gel (Thermo Fisher Scientific), transferred to a nitrocellulose membrane (GE Healthcare) and visualized using a Phosphoimager (Fujifilm). After cutting the correct region from the nitrocellulose membrane using a cutting mask, RNA was isolated by Phenol/Chloroform/Isoamylalcohol (pH = 8.0, Sigma Aldrich) and the mix was transferred to a 2 ml Phase Lock Gel Heavy tube (Quantabio). After separation of the phases the RNA was precipitated by the addition of 0.75 μl GlycoBlueTM Coprecipitant (ThermoFisher), 40 μl 3 M sodium acetate pH 5.5 and 1 ml 100% ethanol. It was mixed and stored at -20 °C for at least two hours. Reverse transcription was performed on the resulting purified RNA using Superscript III (Thermo Fisher Scientific) according to manufacturer’s instructions. Then cDNA was immobilized on MyONE SILANE beads (Thermo Fisher Scientific) as described by the manufacturer and the second adapter was ligated. The library was amplified for six PCR cycles with short primers followed by a ProNex size selection step (Promega) in a 1:2.95 (v/v) sample/bead ratio according to manufacturer’s instructions. The size-selected library was amplified for 13/16 cycles (endogenous PURA, anti-PURA^12D11^, HeLa cells; sample 1 & 4 and sample 2 & 3, respectively), 22 cycles (endogenous PURA, anti-PURA^12D11^, NPCs), 12 cycles (overexpressed FLAG-PURA, anti-PURA^12D11^, HeLa cells), 13/14 cycles (overexpressed FLAG-PURA, anti-FLAG, HeLa cells; sample 1/2, respectively) using Illumina primers (**Supplementary Table S9**). Afterwards, a second size selection was performed using ProNex chemistry (Promega) at 1/2.4 (v/v) sample/bead ratio according to manufacturer’s instructions and cDNA was eluted in 20 μl MQ water. The final library was analyzed using a D1000 Chip (Agilent) in a Bioanalyzer (Agilent). Samples were equimolarly mixed before sequencing on an Illumina HiSeq1500 platform with 75-110-nt single-end reads and 6.7-29.5 million reads per sample (**Supplementary Table S1**).

### Processing of iCLIP reads

Basic quality controls were done with FastQC (v0.11.8) (https://www.bioinformatics.babraham.ac.uk/projects/fastqc/). The first 15 nt of iCLIP reads hold a 6 nt sample barcode and 5+4 nt unique molecular identifiers (UMIs) flanking the sample barcode. Based on the sequence qualities (Phred score) in this 15 nt barcode region, reads were filtered using the FASTX-Toolkit (v0.0.14) (http://hannonlab.cshl.edu/fastx_toolkit/) and seqtk (v1.3) (https://github.com/lh3/seqtk/). The sample barcodes, found on positions 6 to 11 of the reads, were used to de-multiplex the set of all quality filtered reads using Flexbar (46) (v3.4.0). Afterwards, barcode regions and adapter sequences were trimmed from read ends using Flexbar, requiring a minimal overlap of 1 nt of read and adapter. UMIs were added to the read names and reads shorter than 15 nt were removed from further analysis. Downstream analysis was done as described in Chapters 3.4 and 4.1 of Busch et al. (47). GENCODE (30) release 31 genome assembly and annotation were used during mapping.

### Binding site definition

Using the procedure described in (47), we defined 5-nt wide binding sites on the iCLIP data for the endogenous PURA (anti-PURA^12D11^) from HeLa cells. In brief, the processed crosslink events of the four biological replicates were merged into two pseudo-replicates and subjected to peak calling by PureCLIP (version 1.3.1) (48) with default parameters. To define binding sites, PureCLIP sites closer than 5 nt were merged into regions, and isolated PureCLIP sites without an adjacent site within 4 nt were discarded. Binding site centers were defined iteratively by the position with the highest number of crosslink events and enlarged by 2 nt on both sides to obtain 5-nt binding sites. Binding site centers were required to harbor the maximum PureCLIP score within the binding site. Furthermore, binding sites were filtered for reproducibility by requiring a binding site to be supported by a sufficient number of crosslink events in at least two out of four replicates. The threshold for sufficient crosslink event coverage, using the 0.05 percentile, was determined as described in (47).

### Genomic location of binding sites and crosslink events

We mapped binding sites or crosslink events to the gene and transcript annotation from GENCODE (release 31, genome version GRCh38.p12) (**Figure 3A, B**) (30). Genes of gene level 3 were included only when no genes of higher level were overlapping. Similarly, transcripts of transcript level NA were included only when no transcripts of levels 1-3 were annotated for the gene. When a binding site overlapped with two genes, one was chosen at random. For the assignment of PURA binding sites to transcript regions (**Figure 3B**), when binding sites overlapped with different regions from different transcripts, the region was chosen by a hierarchical rule with 3’UTR > 5’UTR > CDS > intron, which we established from visual evaluation of the crosslink event and binding site distribution. We divided the PURA-bound mRNAs into groups according to the location of the strongest PURA binding site on the transcript (3’UTR, n = 2,462; 5’UTR n = 98; CDS, n = 1,646; intron, n = 1; noncoding RNA = 185). To normalize the number of binding sites per region to the lengths of the respective regions (**Supplementary Figure S6H**), we selected all genes with at least one PURA crosslink in any region and calculated a mean region length over all transcripts annotated for a gene. We then summed up the mean region lengths of all selected genes and divided the number of binding sites per region by this sum.

**Figure 3.**
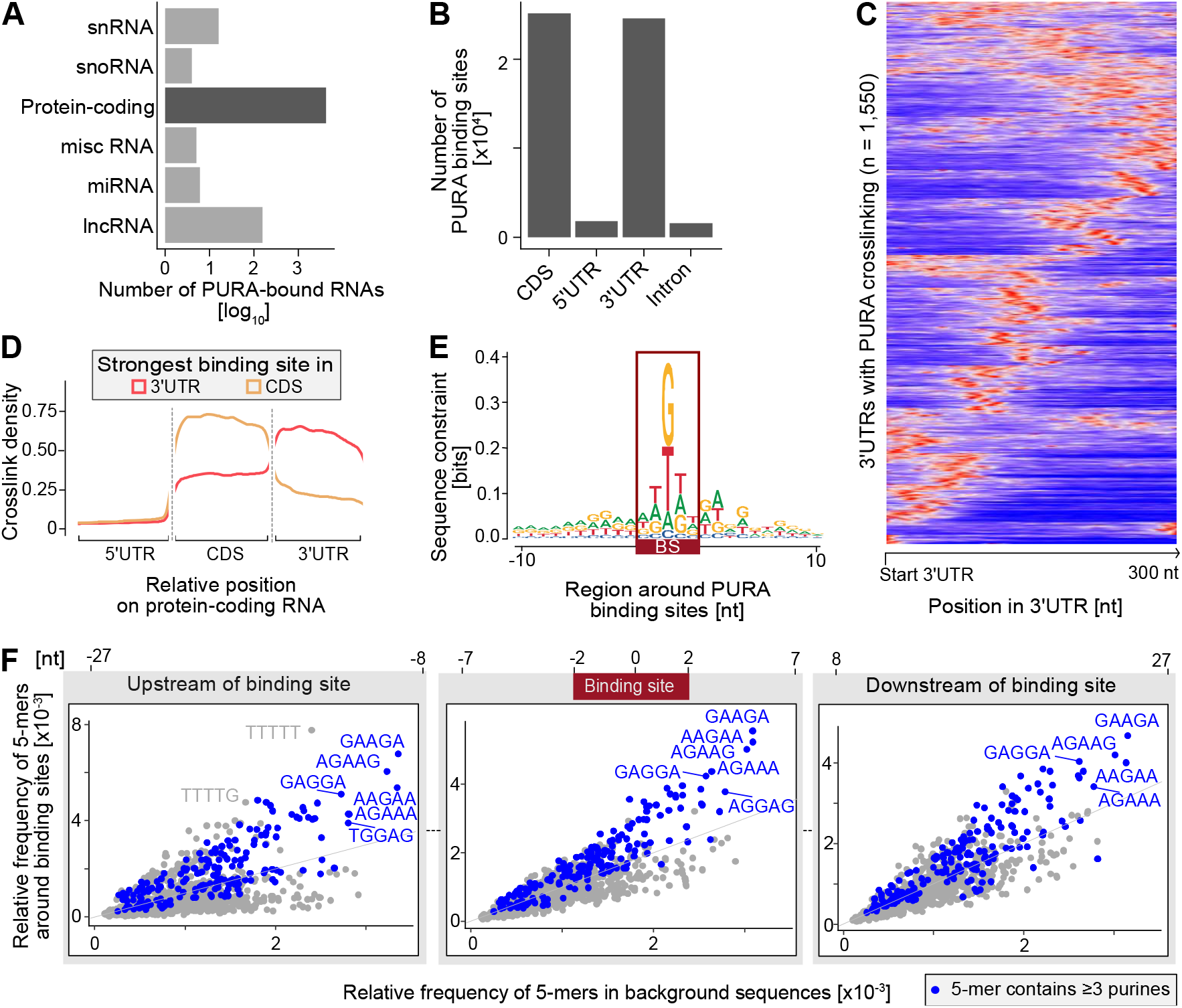
PURA binds to CDS and 3’UTRs of mRNAs. **(A)** Transcript types bound by endogenous PURA in HeLa cells show its preference for binding to protein-coding RNAs. Bars indicate the number of bound RNAs of each type on a log_10_ scale. **(B)** PURA binding sites per transcript region in protein-coding transcripts. Most binding sites are located in 3’UTRs or CDS. Bars show total number of binding sites per region. **(C)** PURA displays one or more distinct peaks in the 3’UTR of bound transcripts. Heatmap shows normalized, smoothed distribution of PURA crosslink events in the first 300 nt of 1,550 exemplary 3’UTRs with intermediate crosslink signal (10^2^-10^6^ crosslink events per window). **(D)** Metaprofile of PURA crosslink events along transcripts with strongest PURA binding site in the 3’UTR (red) or CDS (beige). Smoothed density of crosslink events is shown against scaled positions within the respective region. **(E)** Sequence logo of the nucleotide composition in a 21-nt window around PURA binding sites reveals purine enrichment. Y-axis shows sequences constraint in bits. **(F)** Comparison of the relative frequency of 5-mers in PURA-bound RNAs (y-axis) and a random background (x-axis) is shown for three locations around the binding site: Left – [-27 nt; -8 nt] upstream of the binding site center, middle – [-7 nt; 7 nt] around the binding site center, right – [8 nt; 27 nt] downstream of the binding site center (see annotation above). 5-mers with three or more purines are colored in blue.

The metaprofiles of the distribution of crosslink events along the transcript regions for mRNAs with strongest binding sites in the 3’UTR or CDS (**Figure 3D**) was calculated and visualized in R using the Bioconductor package cliProfiler (http://bioconductor.org/packages/release/bioc/html/cliProfiler.html).

### Comparison of crosslink patterns

We compared the distribution of PURA crosslink events from the iCLIP data for endogenous PURA (anti-PURA^12D11^) from HeLa cells to the two iCLIP datasets in which FLAG-tagged PURA was overexpressed in HeLa cells and immunoprecipitated with either anti-PURA^12D11^ or an anti-FLAG antibody using the following methods: First, to visualize the crosslink patterns in 3’UTRs, we generated heatmaps of the first 300 nt of 1,550 3’UTRs or 573 CDS with intermediated PURA crosslink coverage (10^2^-10^6^ crosslink events per window). In order to account for expression level and other differences, the crosslink events in each window were scaled to the minimum (set to 0) and maximum (set to 1) therein (“min-max normalization”) followed by spline-smoothing using the smooth.spline function (R stats package version 4.1.1, spar = 0.5) and inflated dimensions (dim = 500). The heatmaps show the min-max normalized, spline-smoothed crosslink patterns for the three iCLIP experiments (**Figure 3C, Supplementary Figure S4B**). Second, we assessed differences in the iCLIP signal using metaprofiles of the min-max normalized signal in a 65-nt window around all PURA binding sites identified for endogenous PURA in HeLa cells (**Supplementary Figure S4C**).

### Motif analysis

As PURA predominantly acts on mature RNAs, all motif analyses were performed on the sequences of spliced transcripts. Binding sites were transferred from genomic to transcriptomic coordinates with the Bioconductor package GenomicFeatures (version 1.45.2) (49) using GENCODE transcript annotation (release 31, genome version GRCh38.p12) (30). 49,602 binding sites could be unambiguously assigned and were used for further analyses. Sequences around binding sites were extracted with Biostrings::getSeq (R package, version 2.61.2) (https://bioconductor.org/packages/release/bioc/html/Biostrings.html). For sequence logos, a 51-nt window centered at the binding site was used and logos are generated with ggseqlogo (R package, version 0.1). 23,734 out of 49,602 (47.8%) were centered on a G and 30,454 out of 49,602 (61.4%) enclosed three or more purines.

To search for enriched motifs of 5-nt length (5-mers) within and around the PURA binding sites, we calculated the frequencies of all overlapping 5-mers with Biostrings::oligonucleotideFrequency (version 2.61.2) (https://bioconductor.org/packages/release/bioc/html/Biostrings.html) in three windows: (i) [-7 nt; 7 nt], i.e., including the binding sites plus 5 nt flanking on either side, and (ii) [-27 nt; -8 nt] and [8 nt; 27 nt], i.e., representing the 20 nt on either side of the binding site-containing windows. For comparison, we randomly selected 49,602 5-nt windows from expressed transcripts (i.e., harboring at least one PURA crosslink event).

### Accessibility prediction

We used RNAplfold (from the ViennaRNA package, version 2.4.17) (50) to predict the accessibility in and around PURA binding sites on transcripts of at least 501 nt length (n = 48,525) in comparison to randomly selected positions (n = 10,017). Binding sites were assigned to transcripts as described above. In brief, we used 501-nt windows either around PURA binding sites or randomly selected and predicted the probability for each single nucleotide to be single-stranded. RNAplfold was set to a sliding window w = 100 nt and a maximum span l = 30 nt. As the obtained unpaired probabilities follow a bimodal distribution, we transferred them to log-odds to obtain a bell-shaped distribution: log(odds ratio) = log(prob/1-prob). These were further converted into z-scores using the mean log-odds ratios of the binding sites versus the mean and standard deviation of the background regions (drawing 1,000 regions for 1,000 times). In **Supplementary Figure S6I**, results are displayed only for the 201 nt in the center of the 501-nt window as RNA folding is strongly biased toward single-strandedness at the edges of a given RNA sequence. *P* values were calculated from the z-scores using 2*pnorm(-abs(z_score) and then adjusted for multiple testing with Benjamini Hochberg correction.

### Comparison to published transcriptomes

We used published processed RNA-seq data comparing RNAs in stress granules (human osteosarcoma U-2 OS cells) or P-bodies (human embryonic kidney HEK293 cells) to total RNA (42,51), taking RNAs with FDR < 0.01 and log_2_-transformed fold change (l2fc) > 0 (as calculated by the cited authors) to belong to the respective P-body or stress granule transcriptome.

To determine the overlap of PURA-bound RNAs with dendritically localized RNAs, we used data from (52) that summarizes eight studies describing the dendritic transcriptome in mouse neurons from enrichment in RNA-seq data of dendrites compared to complete neurons. We used only RNAs that were matched to their human orthologs using the Ensembl database (version GRCh38.p13) (53) keeping only one-to-one orthologs (n = 5,070). From these, only RNAs expressed in HeLa cells (baseMean > 0 in our RNA-seq data) were considered for the overlap (n = 4,522). From the dendritically localized RNAs, we only considered RNAs that were found enriched in at least three of the eight studies from (52) (n = 337).

Significance of overlaps was tested using Fisher’s exact test, and differences in fold change distributions between groups were tested using two-sided Wilcoxon Rank-sum test.

### Functional enrichment analyses

We performed functional enrichment analysis in R using the “hypergeometric” mode of the hypeR package (version 1.9.1) on the gene sets from REACTOME (version 7.5.1) and Gene Ontology (version 7.5.1) databases (54,55). HypeR adjusts the gene sets connected to every term specifically to genes present in a given background set. We used the following background sets for the enrichment analyses: All genes with at least one PURA crosslinked nucleotide as background for enrichment of genes bound by PURA (**Figure 2E, Supplementary Figure S6C, D**), all genes with a TPM > 0 in RNA-seq experiment as background for enrichment of RNA expression changes in *PURA* KD (**Supplementary Figure S8B, C**) and all proteins detected by MaxQuant from shotgun proteomics as background for enrichment of protein expression changes in *PURA* KD (**Supplementary Figure S8D, E**). We calculated the gene ratios as the number of detected genes in the respective experiment that belong to a given term divided by the number of genes from the background set of the term. *P* values obtained from HypeR were adjusted using Benjamini Hochberg adjustment and are referred to as FDR. We display the 25 terms with the highest FDR in **Supplementary Figure S6C, D** and **Supplementary Figure S8B-E**. A full list with the statistics of all terms can be found in **Supplementary Table S3, S6, S7**.

## Results

### PURA localizes to the cytoplasm in multiple cell lines

In previous reports, PURA has been associated with both nuclear and cytoplasmic functions. To investigate the subcellular localization of PURA in different cell lines, we raised a monoclonal antibody against the human PURA protein. In the light of earlier studies in which antibodies cross-reacted between PURA and its close paralog PURB (18), we selected as epitope a 21 amino acid (aa) peptide in the unstructured linker region between PUR repeats I and II. This sequence harbors multiple substitutions between PURA and PURB (**Supplementary Figure S1A, B**) but is 100% identical to the mouse Pura ortholog. We confirmed in immunofluorescence (IF) stainings of HeLa cells and Western blots of cell lysates that the obtained antibody (anti-PURA^12D11^) specifically recognized the endogenous PURA protein, detected reduced levels upon *PURA* KD and showed no crossreactivity against recombinant PURB in *E. coli* lysates (**Supplementary Figure S1C-G**). Hence, we used the in-house antibody for the analyses in this study.

Using anti-PURA^12D11^ in IF stainings, we assessed the localization of the endogenous PURA protein in five immortalized cell lines from a diverse set of tissue origins including epithelial, neuronal, kidney, lung and liver tissue. We found that in all cell lines, PURA located predominantly in the cytoplasm (**Figure 1A**). This could be further validated in nuclear-cytoplasmic fractionation experiments in HeLa cells (**Figure 1B, C**). These observations suggest that PURA may be primarily involved in cytoplasmic regulatory processes which we decided to elucidate further.

### PURA acts as a global cytoplasmic RNA-binding protein

Since PURA was previously found to bind RNA *in vitro* (24), we next investigated whether PURA directly associates with RNAs in cells. We therefore performed individual-nucleotide resolution UV crosslinking and immunoprecipitation (iCLIP) experiments, which map the RNA binding of an RBP in living cells (45,56). Endogenous PURA was crosslinked to its bound RNAs in HeLa cells using UV radiation (254 nm) and subsequently immunoprecipitated with anti-PURA^12D11^ (**Figure 2A**). In total, we obtained more than 20 million unique PURA crosslink events from four biological replicates (**Supplementary Table S1**).

Assessing the global distribution of crosslink events before binding site definition, we observed that only a small fraction of PURA crosslink events occurred within introns (**Supplementary Figure S2A**). The relative depletion of introns in the iCLIP experiments suggests that PURA predominantly binds mature RNAs, consistent with its subcellular localization (**Figure 1**) and cytoplasmic mRNA binding.

We next defined PURA binding sites from the iCLIP crosslink events by peak calling and merging adjacent peaks into binding sites of 5-nt width. In total, we identified 55,031 PURA binding sites in the transcripts of 4,391 genes (**Figure 2B, C, Supplementary Table S2**). The strength of PURA binding at these sites, measured as the relative number of crosslink events inside the binding site, was highly reproducible between replicates (**Supplementary Figure S2B**).

To validate PURA binding to RNAs by orthogonal methods, we performed electrophoretic mobility shift assays (EMSA) on 8 selected PURA-bound RNAs. We thereby confirmed direct PURA binding to the iCLIP-identified RNA binding sites in *STARD7, COX7C, NEAT1, LSM14A, NDUFS5, CLTA, CLTC*, and *EEF2* (**Figure 2D, Supplementary Figure S3**). As controls, we used the unspecific yeast RNA *ASH1* E3 as well as the previously described PURA target sequence *MF0677* (**Supplementary Figure S3L-N**) (23).

In a control experiment, we overexpressed FLAG-tagged PURA in HeLa cells and confirmed that comparable RNA binding profiles were obtained from immunoprecipitation with either anti-PURA^12D11^ or an anti-FLAG antibody (**Supplementary Figure S4, Supplementary Table S1, Supplementary Material S1**), underlining the specificity of our in-house antibody. However, in addition to the defined binding sites, the PURA overexpression led to increased crosslink events along transcripts in both experiments, indicating unspecific RNA binding by the unphysiologically abundant ectopic PURA. We therefore decided to focus on the RNA binding profiles detected for the endogenous PURA protein.

PURA was repeatedly found as a component of neural transport granules (16-20). Of note, even though HeLa cells are of non-neural origin, the proteins encoded by the PURA-bound RNAs were enriched in cellular compartments of the neuron, axon and dendrite, among others (**Figure 2E**). Also, more than half of previously reported dendritically localized RNAs (52) were bound by PURA in our iCLIP data (170 out of 337, 50.4%; **Supplementary Figure S5A-C**). To directly test RNA binding of PURA in neural cells, we performed an iCLIP experiment for endogenous PURA (anti-PURA^12D11^) in neural progenitor cells (NPCs) derived from human induced pluripotent stem cells. The NPC state was monitored by the rosette morphology of the cells and the expression of several known marker genes for NPC differentiation (**Supplementary Figure S5D, E**). Although we obtained an iCLIP dataset with only a limited signal depth due to the scarce material, the crosslink events for endogenous PURA in NPCs enriched within the PURA binding sites identified from HeLa cells, indicating a similar RNA binding behavior in both cell types (**Figure 2C, F, Supplementary Figure S5F**).

Altogether, we identified more than 50,000 binding sites of endogenous PURA in HeLa cells. The observed RNA binding pattern was independent of the antibody used and comparable between HeLa cells and NPCs. Hence, these results indicate that the binding sites identified in HeLa cells reflect the *bona fide* binding of PURA in living cells.

### PURA binds in coding sequences and 3’UTRs

Examining the distribution of PURA binding, we found that 16% of all expressed transcripts (transcripts per million [TPM] > 0) harbored at least one PURA binding site (median 7 binding sites per transcript; **Supplementary Figure S6A**). The number of detected PURA binding sites moderately increased with the transcripts’ expression levels, most likely reflecting the better representation of more abundant RNAs in an iCLIP experiment (57) (**Supplementary Figure S6B**). Among the highly expressed transcripts (TPM > 10), 46% harbored at least one PURA binding site, underlining the widespread binding of PURA throughout the transcriptome (**Supplementary Figure S6C, D**).

The 4,391 PURA-bound transcripts comprised predominantly protein-coding mRNAs (n = 4,206, 95.5%, **Figure 3A**). These encoded for functions in nervous system development, mitochondrial processes, the innate immune system and RNA metabolism, among others (**Supplementary Figure S6E, F, Supplementary Table S3**). PURA also showed binding to 154 long non-coding RNAs (lncRNAs; **Figure 3A**), including the previously reported PURA target lncRNAs *RN7SL1* (20,58), which has been implicated in transcriptional control, as well as *NEAT1* and *MALAT1*, two architectural RNAs forming distinct nuclear bodies (59) (**Supplementary Figure S6G**). This supports that a minor fraction of PURA localizes to the nucleus and interacts with nuclear RNAs.

Within the protein-coding transcripts, the PURA binding sites were almost equally distributed between the coding sequences (CDS, n = 25,113) and the 3’ untranslated regions (3’UTRs, n = 24,553; **Figure 3B-D, Supplementary Figure S6H**), which together accounted for 93.6% of all binding sites. In contrast, 5’UTRs and intronic regions were almost completely devoid of PURA binding sites. Within the 3’UTRs or CDS, PURA usually displayed one or more defined peaks of binding, as seen in *STARD7* mRNA (**Figure 2C, 3D, Supplementary Figure S4B**).

The transcriptome-wide profiles enabled us to revisit the RNA sequence and structure preferences of PURA. We therefore predicted the local folding probability around PURA binding sites. This suggested a propensity of PURA’s RNA binding sites to be single-stranded (**Supplementary Figure S6I**), contrasting past reports about its double-stranded nucleic acid binding preference and unwinding ability (23,60). The increased single-strandedness coincided with an apparent sequence preference. Beyond the increased uridine content, likely reflecting the known UV-crosslinking bias (61), we observed a strong enrichment for purines (A and G) within and around the PURA binding sites: 47.8% of binding sites were centered on a guanine and 61.4% enclosed three or more purines (**Figure 3E, Supplementary Figure S6J**). Globally, the binding sites and their surrounding sequences were enriched for *k*-mers comprising purine combinations like GAAGA and AAGAA (**Figure 3F**). Together, these observations underline the specificity of PURA to bind single-stranded, purine-rich regions, primarily in CDS and 3’UTRs.

### Loss of PURA causes widespread changes in gene and protein expression

While iCLIP data revealed the RNAs directly bound by PURA, they do not inform about the regulatory effects of binding events on such transcripts. In order to globally determine the impact of PURA on gene expression, we depleted *PURA* in HeLa cells using siRNAs (**Supplementary Figures S1C-G, S7**). This partial loss of PURA mimics the haploinsufficiency in PURA Syndrome patients that harbor heterozygous mutations in the *PURA* gene. By performing transcriptomic analyses, we observed widespread changes in transcript abundance upon *PURA* KD, relative to control HeLa cells: In total, 3,415 transcripts were significantly down- or upregulated upon *PURA* KD (false discovery rate [FDR] < 0.01, **Figure 4A, Supplementary Table S4**).

**Figure 4.**
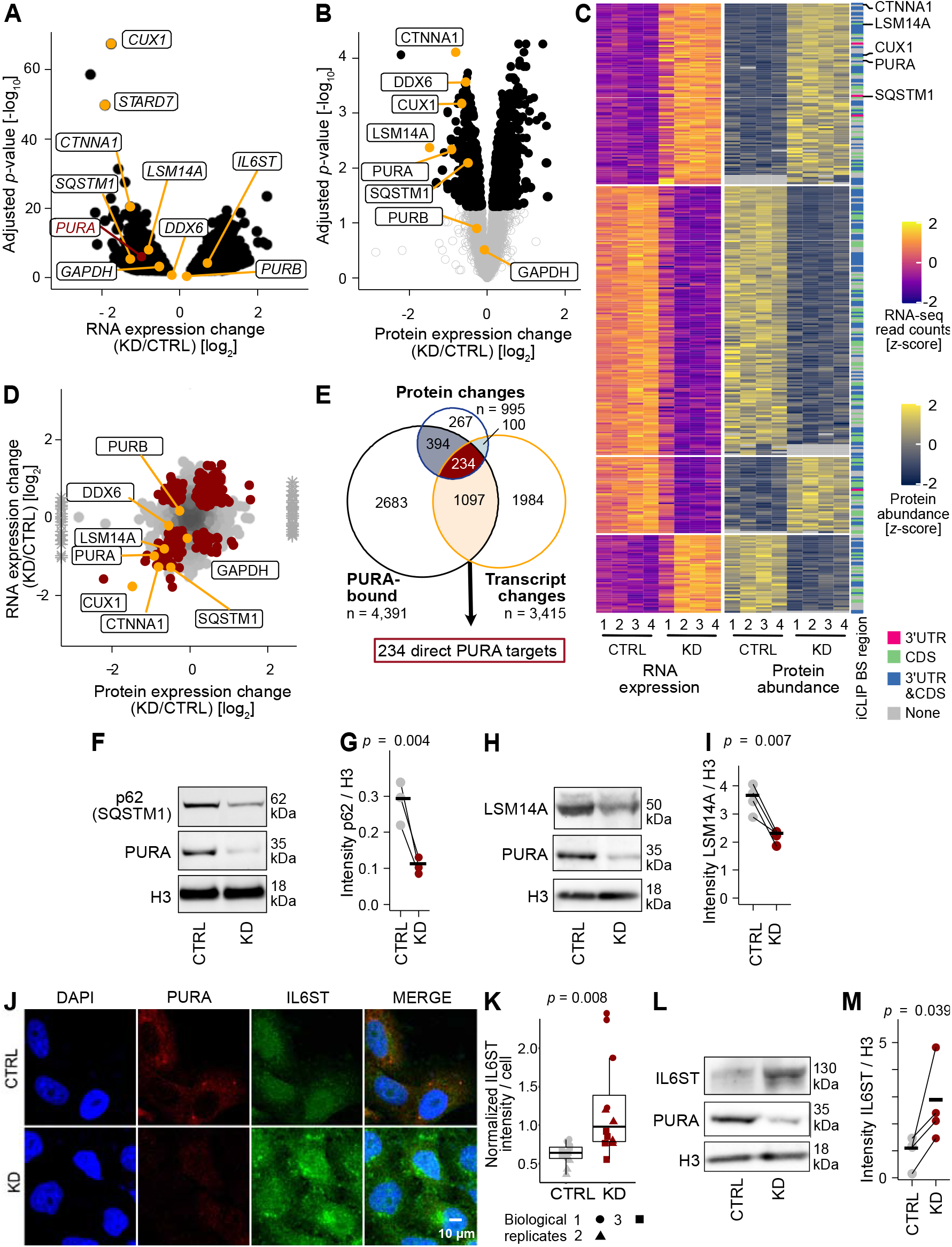
Loss of PURA results in widespread changes in gene and protein expression. **(A)** Differentially expressed transcripts in PURA knockdown (KD) vs. control (CTRL) conditions (n = 4) are shown in a volcano plot giving the RNA expression change [log_2_] against the significance as adjusted P value [log_10_] (black: 3,415 transcripts with FDR < 0.01). **(B)** Changes in protein abundance upon PURA KD (n = 4) are shown in a volcano plot giving the protein expression change [log_2_] against the significance as adjusted P value [log_10_] (black: 995 proteins with FDR < 0.05). **(C)** 334 targets are regulated by PURA on the RNA and protein levels. Heatmap shows z-score-normalized RNA counts and protein abundances of all replicates with the associated PURA-bound transcript region (iCLIP). **(D)** Changes in protein abundances are similar to changes in RNA levels. Scatter plot compares changes in RNA (y-axis) and protein abundance (x-axis) for 4,391 transcripts that were measured in both assays. Targets with significant changes in both data sets (n = 334) are marked in red. Infinite log_2_ fold change values from protein measurements are shown as asterisks. **(E)** Overlap of PURA-bound RNAs with RNAs or proteins significantly changing upon PURA KD. The 234 RNAs overlapping between the three experiments were defined as highly validated PURA targets. **(F-I)** SQSTM1 and LSM14A levels are significantly reduced upon PURA KD. Representative Western blots (F, H) and quantifications of replicates (G, I) are shown in CTRL and PURA KD conditions. PURA KD was confirmed by detection with anti-PURA^12D11^. H3 served as loading controls (n = 3 [SQSTM1] or 4 [LSM14A, IL6ST] biological replicates, paired one-sided Student’s t-test). **(J)** Immunofluorescence staining of PURA (red) together with the IL6ST (green) and DAPI (blue) as a nuclear stain in CTRL (top) and PURA KD (bottom) HeLa cells. Scale bars, 10 μm. **(K)** Quantification (ImageJ) of IL6ST intensity of approximately 80 cells per replicate and condition (3 biological replicates with 4 technical replicates each, unpaired two-sided Student’s t-test). **(L, M)** IL6ST levels significantly increase upon PURA KD. Representative Western blot (L) and quantification of replicates (M) are shown in CTRL and PURA KD conditions. PURA KD was confirmed by detection with anti-PURA^12D11^. H3 served as loading controls (n = 4 biological replicates, paired one-sided Student’s t-test).

To test for changes at the protein level, we performed mass-spectrometric analysis of protein extracts from *PURA* KD and control HeLa cells. We detected a total of 4,351 proteins with good confidence (peptide spectrum match FDR < 1% and protein FDR < 5%) and observed a significant regulation of 995 proteins upon *PURA* KD (FDR < 0.05, **Figure 4B, Supplementary Table S5**). Of these, 334 proteins were significantly regulated also at the transcript level, with the vast majority changing in the same direction in both RNA expression and protein abundance (249, 74.6%, **Figure 4C, D, Supplementary Figure S8A**).

Gene ontology (GO) analysis revealed that the regulated transcripts encoded for diverse functional categories, including mitochondrial functions, vesicle-mediated transport, membrane trafficking and RNA metabolism (**Supplementary Figure S8B, C, Supplementary Table S6**). Similarly, the regulated proteins were enriched in mitochondrial and signaling functions and several membrane and granule components (**Supplementary Figure S8D, E, Supplementary Table S7**).

Out of 334 proteins whose levels changed at both transcript and protein level in response to *PURA* KD, 234 were independently identified as PURA-bound RNAs by iCLIP (**Figure 4E**). These showed similar changes in RNA and protein expression upon *PURA* KD, irrespective of the location of PURA binding sites in the CDS or 3’UTR (**Figure 4C, Supplementary Figure S9A-F**).

The most significantly downregulated transcript was the mRNA encoding the transcription factor CUX1 (**Figure 4A-D**), which is involved in the control of neuronal differentiation by specifically regulating dendrite development, branching, and spine formation in cortical neurons (62). Another strongly regulated PURA target was the cytoskeleton associated Catenin alpha-1 (CTNNA1) that links actin and adherence junction components (63). Furthermore, a coherent downregulation upon *PURA* KD was also observed for the PURA-bound RNAs encoding for the autophagy receptor Sequestome 1 (SQSTM1; also known as p62), consistent with a previous report (64), and the essential P-body component LSM14A. The downregulation of the SQSTM1 and LSM14A proteins was orthogonally confirmed by Western blot analysis (**Figure 4F-I**).

Due to the very different detection limits of RNA-seq and shotgun proteomics, it is likely that we did not detect all direct PURA targets in both data sets. To assess a representative target that did not pass the threshold for a reliable quantification in our proteomics data, we analyzed the levels of interleukin-6 cytokine family signal transducer (IL6ST) using fluorescence microscopy (**Figure 4J, K, Supplementary Figure S9G**) and Western blot (**Figure 4L, M**). As suggested in our RNA-seq data, we observed an increased signal intensity and accumulation of IL6ST in the perinuclear region upon *PURA* KD in both HeLa and normal human dermal fibroblast (NHDF) cells, supporting a role of PURA in IL6ST protein expression.

In essence, we find that loss of PURA results in widespread changes in gene and protein expression, affecting central cellular features such as mitochondria, autophagy and granule assembly. The quantitative comparison of transcriptome and proteome suggests that the majority of changes might originate at the RNA level. Ultimately, we identify a set of 234 targets that change consistently on RNA and protein level upon *PURA* KD and are bound by PURA on the RNA level. As the example of IL6ST shows, the list of PURA targets could potentially be extended beyond this stringent set of targets that are coherently identified by iCLIP, RNA-seq and proteomics.

### PURA binds to P-body transcripts and localizes to P-bodies

Under unstressed conditions, we observed the PURA immunofluorescence signal in cytoplasmic foci (**Figure 1**). To obtain a first indication what kind of foci these might be, we overlapped the PURA-bound RNAs with transcripts that were enriched in P-bodies (42) or stress granules (51). In line with the previously reported PURA localization to stress granules (21), more than half of the stress granule-enriched transcripts were bound by PURA (44%, *P* value < 2.2×10^−16^, Fisher’s exact test; **Figure 5A, Supplementary Figure S10A-C**). Surprisingly, however, we detected an almost equally strong overlap with transcripts enriched in P-bodies (42) (43%, *P* value < 2.2×10^−16^, Fisher’s exact test; **Figure 5A, Supplementary Figure S10D-F**), indicating that PURA might be linked to these granules as well.

**Figure 5.**
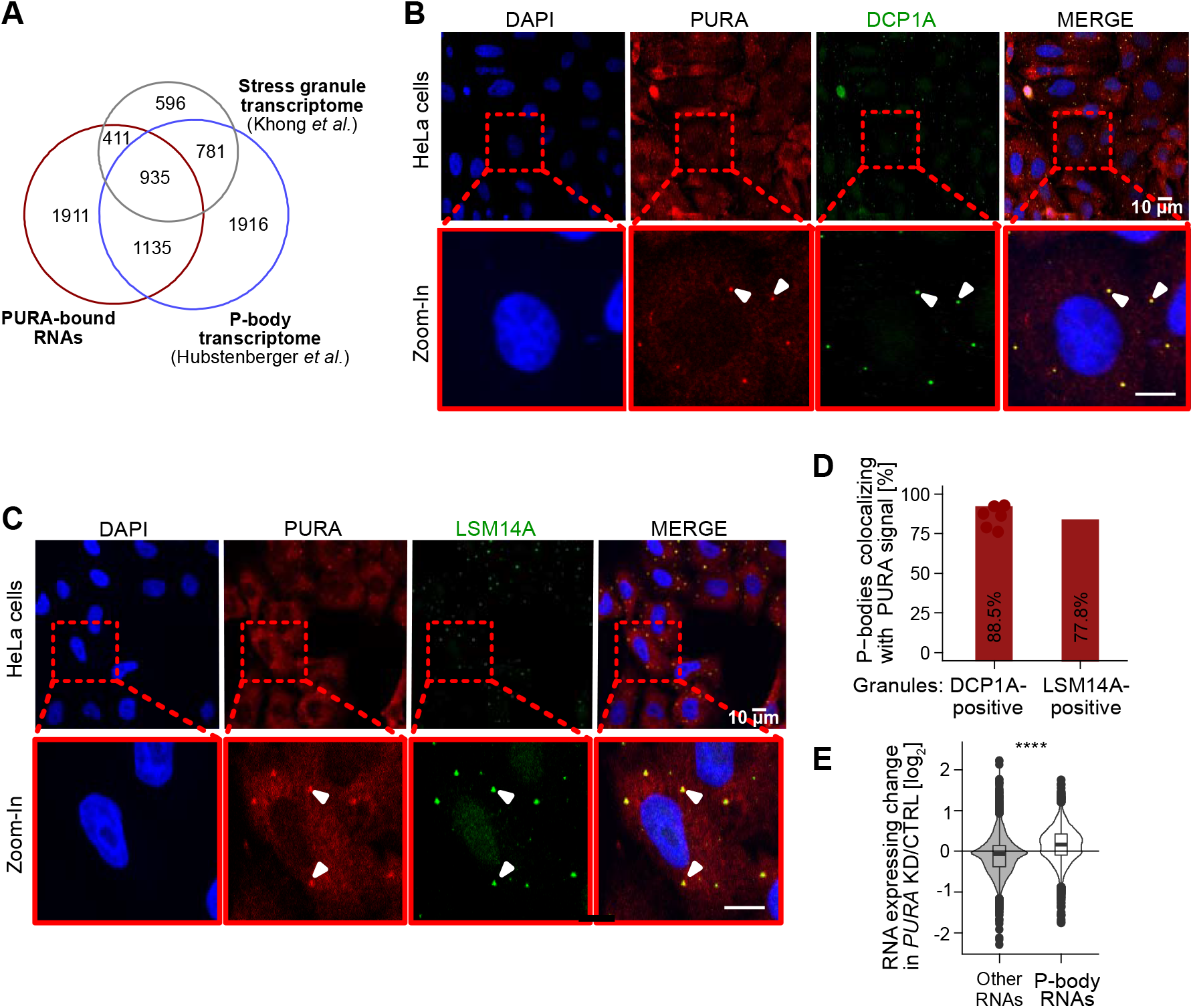
PURA localizes to P-bodies and P-body RNAs are upregulated upon PURA KD. **(A)** Venn diagram depicts overlap of PURA-bound RNAs (red) with the transcriptomes of P-bodies (blue, n = 4,767, odds ratio 5.7 [5.3, 6.2] (95% CI), P value < 2.2×10^−16^, Fisher’s exact test) (42) and stress granules (grey, n = 2,723, odds ratio 12.0 [10.8, 13.4], P value < 2.2×10^−16^, Fisher’s exact test) (51). **(B)** Confocal micrographs of PURA (red), P-body marker DCP1A (green) and colocalized signal (yellow). Nuclei were stained with DAPI (blue). Arrowheads indicate examples of P-bodies with overlapping staining. Scale bars, 10 μm. **(C)** Staining as in (A) using LSM14A as P-body marker (measured in 555 nm channel, depicted in green). Colocalization (yellow) of PURA (red) and LSM14A (green). Scale bars, 10 μm. **(D)** Colocalization of PURA (anti-PURA^12D11^) with DCP1A- and LSM14A-positive granules (6 samples each, mean over all samples is given by bars). Granules were defined as described in Methods. **(E)** P-body RNAs are upregulated in PURA KD. Violin plot of RNA expression changes (log_2_ fold change, PURA KD/CTRL) for RNAs enriched in P-bodies (blue) versus all other RNAs (grey) (P value < 0.0001, unpaired two-sided Wilcoxon Rank-sum test).

To test for a direct association, we used immunofluorescence staining to overlay PURA with the P-body marker proteins DCP1A and LSM14A. Indeed, we identified a strong colocalization of PURA with both marker proteins in HeLa and NHDF cells (**Figure 5B, C, Supplementary Figure S10G, H**). Of note, we detected PURA in 88.5% of all DCP1A-positive granules and 77.8% of all LSM14A-positive granules (**Figure 5D**). This indicates that under basal growth conditions, PURA efficiently localizes to P-bodies in HeLa and NHDF cells.

### PURA depletion reduces the number of P-bodies per cell

Based on the strong association of PURA with P-bodies, we wondered whether PURA binding directly affects the abundance of the P-body-associated RNAs. Intriguingly, the P-body-enriched transcripts were selectively upregulated upon *PURA* KD, while the remaining RNAs were not affected (**Figure 5E**). Based on these observations, we hypothesized that P-bodies might be altered in the absence of PURA.

In order to focus on PURA’s role in the physiological context of unstressed cells, we inspected the PURA iCLIP data from HeLa cells and were intrigued to find that PURA directly bound to *LSM14A* and *DDX6* mRNAs, encoding two P-body core components (**Figure 6A, B**). Moreover, LSM14A was strongly downregulated upon *PURA* KD at the transcript as well as protein level, and DDX6 was downregulated at the protein level (**Figure 4A-D**). This was orthogonally validated by a significant reduction in LSM14A and DDX6 protein in Western blot analysis after *PURA* KD, relative to control HeLa cells (**Figure 4H, I, Supplementary Figure S11A, B**).

**Figure 6.**
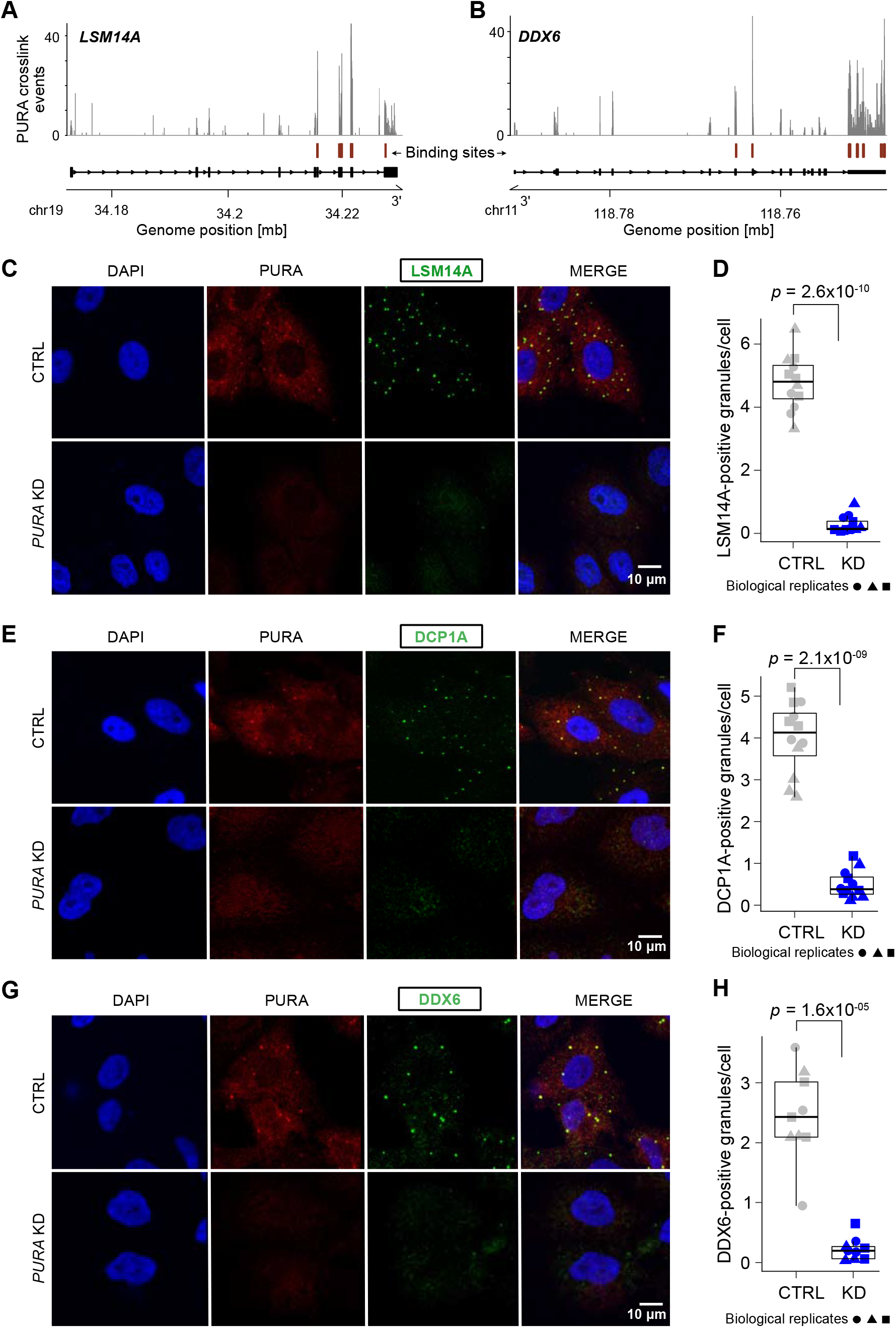
PURA KD leads to a loss of P-bodies. **(A, B)** Genome browser view shows the pileup of crosslink events of endogenous PURA from HeLa cells at binding sites (red) in the LSM14A (A) and DDX6 (B) transcripts. **(C-H)** Left: Immunofluorescence staining of PURA (red) together with the P-body proteins LSM14A (C), DCP1A (E) and DDX6 (G) (measured in 555 nm channel, depicted in green) and DAPI (blue) as a nuclear stain in CTRL (top) and PURA KD (bottom) HeLa cells. Scale bars, 10 μm. Right: Quantification (ImageJ) of LSM14A-positive granules on approximately 400 cells per replicate (n = 3) comparing CTRL and PURA KD conditions. Shown is the number of granules positive for LSM14A (D), DCP1A (F) and DDX6 (H) per cell for all replicates. 3 biological replicates (independent cultures) are indicated by different icons, with each containing 4 technical replicates (multiple images). P value from unpaired two-sided Student’s t-test on all replicates.

Strikingly, when analyzing either LSM14A-, DCP1A- or DDX6-positive granules in *PURA* KD cells using IF staining, we observed a drastic decrease in granule numbers per cell using all markers (**Figure 6C-H**). Although several proteins modulate the appearance of P-bodies, LSM14A and DDX6 are two of a handful of currently known factors that have been shown to be essential for P-body formation (41). To test whether the PURA-dependent reduction of LSM14A and DDX6 globally impairs P-bodies, we visualized P-body occurrence using IF staining for the P-body marker DCP1A. Remarkably, *PURA* KD led to a dramatic reduction in P-body number in HeLa and NHDF cells (**Figure 6C-H, Supplementary Figure S11E-H**). Importantly, in contrast to LSM14A and DDX6 (**Figure 4H, I, Supplementary Figure S11A, B**), the protein levels of the P-body marker DCP1A remained unchanged upon *PURA* KD (**Supplementary Figure S11C, D**), suggesting that PURA acts selectively by modulating the LSM14A and DDX6 levels in tested cell lines. These observations were validated using a second siRNA against *PURA* (**Supplementary Figure S11I, J**).

Together, our data suggest that PURA binds the *LSM14A* and *DDX6* mRNAs and regulates the concomitant protein levels. It is conceivable that the combined reduction of both factors causes the almost complete loss of P-bodies upon *PURA* KD. Given the important role of P-bodies in cellular RNA surveillance, our findings reveal a putative link between the neuronal PURA Syndrome and the cytoplasmic control of mRNA levels and P-body condensation.

## Discussion

### PURA is a cytoplasmic RNA-binding protein

The PURA protein was initially described as a transcription factor with binding preferences for purine-rich single-stranded DNA regions (13,14). Consistent with a nuclear role of PURA, we do observe binding of PURA to nuclear RNAs such as *NEAT1* and *MALAT1*. The minor fraction of PURA in the nucleus might be involved in transcriptional regulation as reported previously (3,15). While this might be one of PURAs functions, it was subsequently shown to associate with cellular RNAs and to bind single-stranded DNA and RNA sequences *in vitro* in the same way and with similar affinities (23). In this study, we developed a paralog-specific antibody to demonstrate that the vast majority of PURA protein resides in the cytoplasm in a range of different cell lines. This is in line with more recent studies describing PURA in cytoplasmic RBP granules (17-19,21). It also underlines the need to study PURA’s function in processes other than DNA regulation.

To date, a physiological RNA binding function of PURA had only been shown for a few specific target RNAs like the lncRNA *RN7SL1* (20,58) or the circular RNAs circSamD4 (65) and circCwc27 (66). Most other studies were based on either *in vitro* binding assays, colocalization studies or enrichment in membrane-less organelles (reviewed in (3)). Here, we provide the first transcriptome-wide analysis of PURA’s RNA binding in living cells. Our iCLIP data show that PURA binds to almost half of all expressed mRNAs, thereby bearing the potential to regulate a multitude of different cellular processes (see below).

In line with the first description of PURA as purine rich element binding protein (13,14,67,68), we identified a binding preference for purine-rich RNA regions. Similar to previous studies (23,24), we did not observe a distinct sequence motif. A degenerate sequence preference is characteristic for many RBPs (69) and might suggest that PURA’s RNA recognition in cells is driven by additional factors, such as RNA secondary structure or co-factor binding.

### PURA regulates mRNAs associated with mitochondria, neural transport granules and autophagy

Omics allow for the global characterization and quantification of pools of biological molecules that translate into a holistic view of cellular functions and dynamics. Our integration of multiscale omics data revealed widespread changes in expression upon loss of PURA in cells. We identify more than 3,000 dysregulated transcripts upon *PURA* KD and almost 1,000 dysregulated proteins. Despite the high sequence similarity to PURA’s close paralog PURB, we do not observe a compensatory upregulation of PURB on RNA or protein level upon *PURA* depletion (**Figure 4A, B, D**), speaking against compensatory effects in the data. The observed changes could result from complementary RNA-regulatory mechanisms, which may include translation, RNA stability or degradation, RNA transport as well as subcellular localization and phase condensation, among others.

Integration with transcriptome-wide RNA binding sites revealed 234 putative direct targets of PURA. The molecular roles of these targets link PURA to various processes including interleukin signaling, autophagy, mitochondrial homeostasis and neurodevelopment. One interesting target that was directly bound by PURA and upregulated upon *PURA* KD is the *IL6ST* mRNA encoding interleukin-6 cytokine family signal transducer (IL6ST; also known as gp130). IL6ST is a signal transducer shared by many cytokines, including interleukin 6, ciliary neurotrophic factor, leukemia inhibitory factor, and oncostatin M (70). Patients with PURA Syndrome appear to have a compromised immune response that may be responsible for the severe infections reported by affected families (personal communication). It is possible that an overexpression of IL6ST upon *PURA* haploinsufficiency may contribute to such symptoms.

Another direct PURA target with potential links to the phenotype of PURA Syndrome patients is the *SQSTM1* mRNA encoding the autophagy receptor Sequestome 1 (SQSTM1; also known as p62). In a zebrafish model for C9 ALS, RNA-triggered toxicity could be rescued through ectopic overexpression of PURA, and this rescue could be related to the upregulation of SQSTM1 by PURA (64). Interestingly, we observed that PURA directly binds to *SQSTM1* mRNA and multiple other targets from the autophagy pathway. This suggests that PURA may act as a modulator of autophagy in human cells, which has long been proposed as a neuroprotective mechanism in cellular surveillance (71).

Furthermore, several of the newly identified PURA targets that are downregulated upon *PURA* KD are involved in mitochondrial functions, including the *STARD7, COX7C* and *FIS1* mRNAs. Indeed, mitochondrial respiration was already described to be impaired upon *PURA* depletion (72). Since mitochondrial dysfunctions are associated with various neuronal disorders (73), it is tempting to speculate that mitochondrial defects caused by *PURA* haploinsufficiency may contribute to the neurodevelopmental phenotype of PURA Syndrome patients. This may converge with direct effects of PURA on neurodevelopmentally relevant targets. In line with this, PURA has been associated with neuronal RNA transport in different studies in mouse and human cells. For instance, a misdistribution of the dendritic marker protein MAP2 was reported in one *PURA* KO mouse model (8). The direct PURA targets identified in our study will be relevant in establishing the neuronal and neurodevelopmental functions of PURA in the future.

### PURA depletion leads to an upregulation of P-body-associated transcripts and a loss of P-bodies

Cytoplasmic RBPs are known to play important roles in the formation of cytoplasmic membrane-less organelles such as P-bodies and stress granules (74). While omics studies provided a broad view on their transcriptomes and proteomes (42,51), only a small number of proteins are considered essential for P-body formation, including DDX6, 4E-T and LSM14A (41).

Here, we show that PURA is enriched in P-bodies and binds to more than 40% of the P-body transcriptome (51). Interestingly, upon *PURA* KD, P-body-enriched transcripts are globally upregulated. This indicates that RNAs stored in P-bodies might be destabilized in these condensates, while they are potentially selectively stabilized upon dissolving of P-bodies. It will require further studies to understand if this observation has functional consequences, for instance for the etiology of patients with PURA Syndrome. In addition, we find that depletion of PURA in different human cell lines leads to a loss of P-bodies. Interestingly, this observation is linked to direct binding of PURA to the *LSM14A* and *DDX6* mRNAs, a downregulation of *LSM14A* mRNA and a concomitant downregulation of the LSM14A and DDX6 proteins. Since both proteins are essential P-body factors, their combined PURA-dependent depletion most likely triggers the observed loss of P-bodies. Interestingly, loss of DDX6 is associated with another neurodevelopmental disorder in which a mutation in the *DDX6* gene leads to several phenotypes including intellectual disability, as observed in PURA Syndrome (75). It would be interesting to understand the similarities in phenotypes between patients with *PURA* and *DDX6* mutations.

P-bodies were initially proposed as a location of RNA depletion (76). Seemingly in line with this, we find that RNAs of the P-body transcriptome are globally upregulated upon *PURA* KD and P-body loss. However, more recently P-bodies were proposed to act in RNA storage rather than depletion (42). Moreover, stress granules and P-bodies were shown to partially share their proteomes and transcriptomes (42) and suggested to dynamically interact with each other (77,78). Since in a previous study (21), PURA was found in stress granules under stressed conditions and we now show its P-body localization, it becomes clear that PURA is one of the proteins shared between both granule types.

### Implications for understanding the etiology of the PURA Syndrome

As PURA Syndrome is described to result from a haploinsufficiency and thereby lowered cellular PURA levels, our findings on PURA-dependent, LSM14A/DDX6-mediated P-body formation could potentially relate to a molecular dysfunction in patients. Altered granule dynamics might play a role in the patho-mechanism of PURA Syndrome. Similarly, the observed changes in the sequestome degradation machinery and the mitochondrial proteome may be in functional correlation to symptoms of PURA Syndrome patients. Among the most downregulated targets was also the transcription factor CUX1 that is involved in the control of neuronal differentiation (62). This and other factors suggest a connection of our molecular observations of misregulated factors to the neurodevelopmental delay of patients.

While the above-mentioned potential links are highly suggestive, it should be considered that the spectrum of symptoms of PURA Syndrome patients is broad and variable (6,7). Furthermore, the understanding of the medical condition of patients has only begun to be investigated, rendering it very difficult to directly and safely correlate molecular PURA-dependent events to symptoms of patients with haploinsufficiency in the *PURA* gene. This consideration is even more important as not all molecular changes are likely to have an impact on the symptoms observed in humans. While this study opened the door for an unbiased and systematic understanding of molecular PURA-dependent pathways, it will require more research on both ends, clinical and non-clinical, to convincingly connect our findings with the symptoms in PURA Syndrome patients.

## Data availability

All sequencing data is available in the Gene Expression Omnibus (GEO) under the SuperSeries accession number GSE193905. The collection includes the RNA-seq data from *PURA* KD and CTRL HeLa cells (GSE193900) as well as the iCLIP data for endogenous PURA (anti-PURA^12D11^ antibody) in HeLa cells (GSE193901) and NPC (GSE193902) as well as for overexpressed FLAG-PURA in HeLa cells precipitated with anti-PURA^12D11^ (GSE193904) and anti-FLAG antibody (GSE193903).

The mass spectrometry proteomics data have been deposited to the ProteomeXchange Consortium via the PRIDE (79) partner repository with the dataset identifier PXD030266.

Scripts used to process the files are accessible under the GitHub repository located at: https://github.com/ZarnackGroup/Publications/blob/main/Molitor_et_al_2022/

## Acknowledgements

The authors would like to thank the members of the Niessing and Zarnack labs for support and discussion. We thank Saskia Hutten for help with immunofluorescence experiments, Annika Niedner-Boblenz and Nadine Körtel for help in setting up iCLIP experiments and Stefan Krebs for help in planning and performing the sequencing runs. We also thank Jana Tretter, Verena Kirchner, Elke Pertler, Deniz Yavru, Hania Ahmed, Mirko Brüggemann, You Zhou, Robert Janowski and Vera Roman for advice and technical assistance. We particularly thank You Zhou, Mirko Brüggemann and Mario Keller for internal code review. We would like to thank Ruth Brack-Werner for providing us with additional cell lines, Robert Schneider for using their BioAnalyzer and the confocal microscopy facility at HMGU for using their microscopes. We also thank Eran Hornstein for helpful comments and discussions.

## Author contributions

L.M. performed iCLIP, *PURA* knockdown, immunofluorescence (IF), Western blot and EMSA, validated the anti-PURA^12D11^ antibody and analyzed data. M.K. performed most bioinformatics analyses of iCLIP, RNA-seq and shotgun proteomics experiments. S.B. performed and quantified IF experiments of PURA and P-body marker proteins and IL6ST as well as Western blots and EMSAs. J.M.-P. prepared, measured and analyzed shotgun proteomics samples. N.S. performed IF experiments for the PURA cell atlas. S.B. performed nuclear-cytoplasmic extraction. C.K. performed Western blot experiments. E.R. differentiated hiPSCs to NPCs and validated NPCs for subsequent iCLIP experiments. D.T. performed Western blot experiments for SQSTM1. A.P. reprogrammed and validated the HMGU12 hiPS cell line. M.P. assisted with IF experiments for P-body marker DCP1A. S.R. performed RNA extraction and library preparation for RNA-seq samples. H.B. planed and supervised sequencing of iCLIP and RNA-seq libraries. A.B. performed initial analysis of iCLIP data sets. S.B. performed Western blots and sequence alignments. The antibody facility headed by R.F. generated the in-house anti-PURA^12D11^ antibody. S.H. planed and supervised shotgun proteomics experiments. M.D. planed and supervised reprogramming and differentiation experiments of hiPSC. P.F.-P. planed and supervised the validation of autophagy marker SQSTM1. J.K. planed and supervised iCLIP experiments. Study was designed by L.M., M.K., K.Z. and D.N.. K.Z. supervised most of the bioinformatics analyses. D.N. supervised most of the experimental work. L.M., M.K., K.Z. and D.N. wrote the manuscript with comments from all co-authors.

