## Supplementary Material for "Depletion of the RNA-binding protein PURA triggers changes in posttranscriptional gene regulation and loss of P-bodies"

* Shared first authors.

Affiliations:

[1] Institute of Structural Biology, Helmholtz Zentrum München - German Research Center for Environmental Health, 85764, Neuherberg, Germany.

[2] Buchmann Institute for Molecular Life Sciences (BMLS) and Faculty Biological Sciences, Goethe University Frankfurt, 60438, Frankfurt, Germany.

[3] Metabolomics and Proteomics Core, Helmholtz Zentrum München - German Research Center for Environmental Health, 85764, Neuherberg, Germany.

[4] Institute of Pharmaceutical Biotechnology, Ulm University, 89081, Ulm, Germany.

[5] iPSC Core Facility, Helmholtz Zentrum München - German Research Center for Environmental Health, 85764, Neuherberg, Germany.

[6] Department of Pediatrics and Adolescent Medicine, Ulm University Medical Center, 89070, Ulm, Germany.

[7] Institute of Molecular Biology (IMB), 55128, Mainz, Germany.

[8] LAFUGA Genomics, Genzentrum, Ludwig-Maximilians University Munich, 81377, Munich, Germany.

[9] Monoclonal Antibody Core Facility, Institute for Diabetes and Obesity, Helmholtz Zentrum München - German Research Center for Environmental Health, 85764, Neuherberg, Germany.

[10] Institute of Stem Cell Research, Helmholtz Zentrum München - German Research Center for Environmental Health, 85764, Neuherberg, Germany.

[11] Division of Drug Discovery and Safety, Leiden Academic Centre for Drug Research (LACDR), Leiden University, 2333 CC Leiden, The Netherlands.

### Content:

**Supplementary Figures ... 4**

Supplementary Figure S1: The monoclonal antibody anti-PURA[12D11] is PURA-specific and discriminates between PURA and PURB. ... 4

Supplementary Figure S2: iCLIP shows global binding of endogenous PURA to RNAs. ... 6

Supplementary Figure S3: EMSAs ... 8

Supplementary Figure S4: Comparison of PURA crosslink patterns from complementary iCLIP experiments validates specificity of the anti-PURA[12D11] antibody. ... 9

Supplementary Figure S5: Validation of results in neural precursor cells (NPCs). ... 11

Supplementary Figure S6: Accessibility and sequence composition at PURA binding sites. ... 13

Supplementary Figure S7: Validation of siRNA-mediated *PURA* knockdown. ... 14

Supplementary Figure S8: Functional enrichment analyses of differentially expressed RNAs and proteins upon *PURA* KD. ... 15

Supplementary Figure S9: The changes in RNA and protein levels upon *PURA* KD are not linked to PURA binding in a particular transcript region. ... 17

Supplementary Figure S10: PURA-bound RNAs are likely to be enriched in the stress granule and p-body transcriptome and PURA localizes in P-bodies in normal human dermal fibroblast (NHDF) cells. ... 20

Supplementary Figure S11: *PURA* KD leads to a loss of P-bodies in NHDF cells as seen in HeLa cells (Figure 6C-H). ... 21

Supplementary Figure S12: uncropped Western blots. ... 24

**Supplementary Material ... 26**

Supplementary Material 1: Comparison of PURA crosslink patterns from complementary iCLIP experiments ... 26

1) iCLIP experiments with overexpressed FLAG-PURA immunoprecipitated with anti-PURA[12D11] and anti-FLAG lead to comparable crosslink patterns, supporting the specificity of the anti-PURA[12D11] antibody. . 26

2) The increase of cellular PURA levels by FLAG-PURA overexpression is accompanied to a higher background signal ... 26

3) PURA binds to similar binding sites in neuronal precursor cells ... 27

**Supplementary Tables ... 28**

Supplementary Table S1: Overview of the four PURA iCLIP experiments. ... 28

Supplementary Table S2: 4,391 PURA-bound genes. ... 28

Supplementary Table S3: Enriched REACTOME and GO cellular component terms for 4,391 PURA-bound genes. ... 28

Supplementary Table S4: Differential RNA expression upon *PURA* knockdown. ... 29

Supplementary Table S5: Differential protein abundance upon *PURA* knockdown. ... 29

Supplementary Table S6: Enriched REACTOME and GO cellular component terms for 3,415 differentially expressed RNAs in *PURA* knockdown. ... 30

### Supplementary Figures

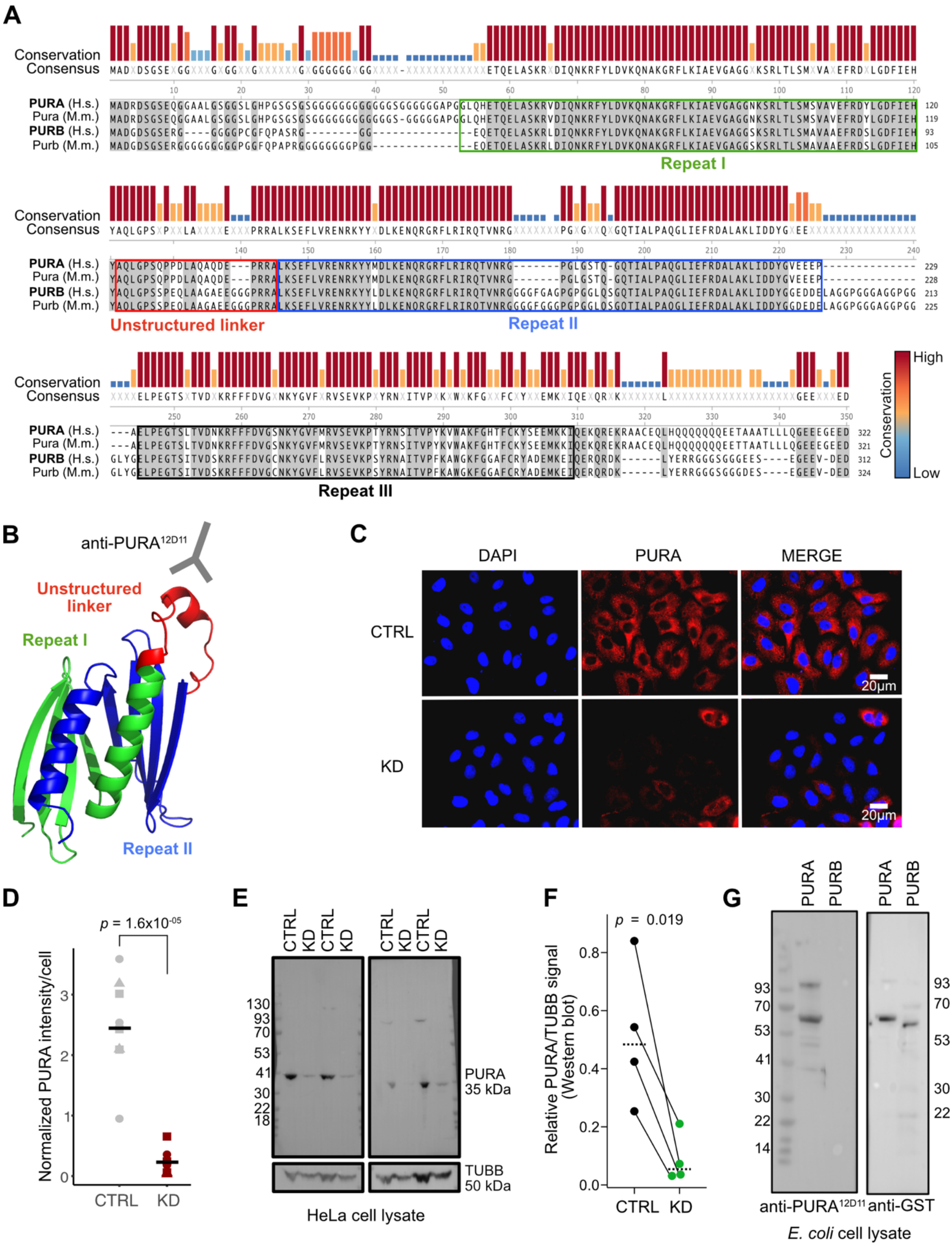

**Supplementary Figure S1: The monoclonal antibody anti-PURA$^{12D11}$ is PURA-specific and discriminates between PURA and PURB.** **(A)** Sequence alignment of the closely related paralogs PURA and PURB from human (*Homo sapiens*, H.s.) as well as Pura and Purb from mouse (*Mus musculus*, M.m.). The 21 amino acid (aa)

peptide from the largely unstructured linker region of PURA (red), used to raise the monoclonal antibody anti-PURA$^{12D11}$, is 100% identical between PURA from human and Pura from mouse, but harbors 6 aa substitutions plus a 3-aa insertion in PURB. Sequence alignment was performed using MEGAX software (1) with MUSCLE algorithm (2). Amino acid sequences were obtained from ensembl.org (3). Marked in red is the unstructured linker region, green and blue are PUR repeats I and II, respectively. Consensus sequence and sequence conservations as colored bars is given on top of the individual sequences (blue – low conservation, yellow – medium conservation, red – high conservation). **(B)** Homology model of human PURA structure as predicted by AlphaFold2 (4) with protein regions colored as in (A). Binding site of anti-PURA$^{12D11}$ antibody to the linker region is additionally indicated. **(C)** Immunofluorescence staining of *PURA* knockdown (KD) and control (CTRL) using PURA$^{12D11}$ antibody (red) and DAPI (blue). Scale bars, 20 µm. **(D)** Quantification of intensity of PURA signal in (C) using ImageJ (n = 16 images), *P* value from two-sided unpaired Student's *t*-test. **(E)** Western blot of endogenous PURA and *PURA* KD in HeLa cell lysates shows depletion of PURA signal and hence specificity of anti-PURA$^{12D11}$. Moreover, the same samples were also used for RNA-seq (see **Figure 4A, Supplementary Figure S7**), further confirming the successful *PURA* KD and the specificity of the PURA$^{12D11}$ antibody. **(F)** Quantification of relative PURA protein amounts in *PURA* KD and CTRL samples from (E) shows a significant PURA downregulation in *PURA* KD samples (paired one-sided Student's *t*-test). Of note, normalization of sample loading with TUBB likely underestimated the observed effect as this protein is itself downregulated by *PURA* KD. **(G)** Western blot towards GST-tagged PURA and PURB recombinantly expressed in *E. coli* lysate (left – anti-PURA$^{12D11}$, right – anti-GST). Anti-PURA$^{12D11}$ specifically detects PURA, while anti-GST detects PURA and PURB. Of note, the specificity of the anti-PURA$^{12D11}$ antibody was further confirmed by subsequent functional assays in this study, such as comparative analyses of iCLIP results performed with over-expressed FLAG-PURA and either anti-FLAG antibody or anti-PURA$^{12D11}$ antibody (**Supplementary Figure S4, Supplementary Table S1**; for a more detailed explanation see **Supplementary Material 1**), which yielded very similar cross-linking patterns. Uncropped images of Western blots are shown in **Supplementary Figure S12**.

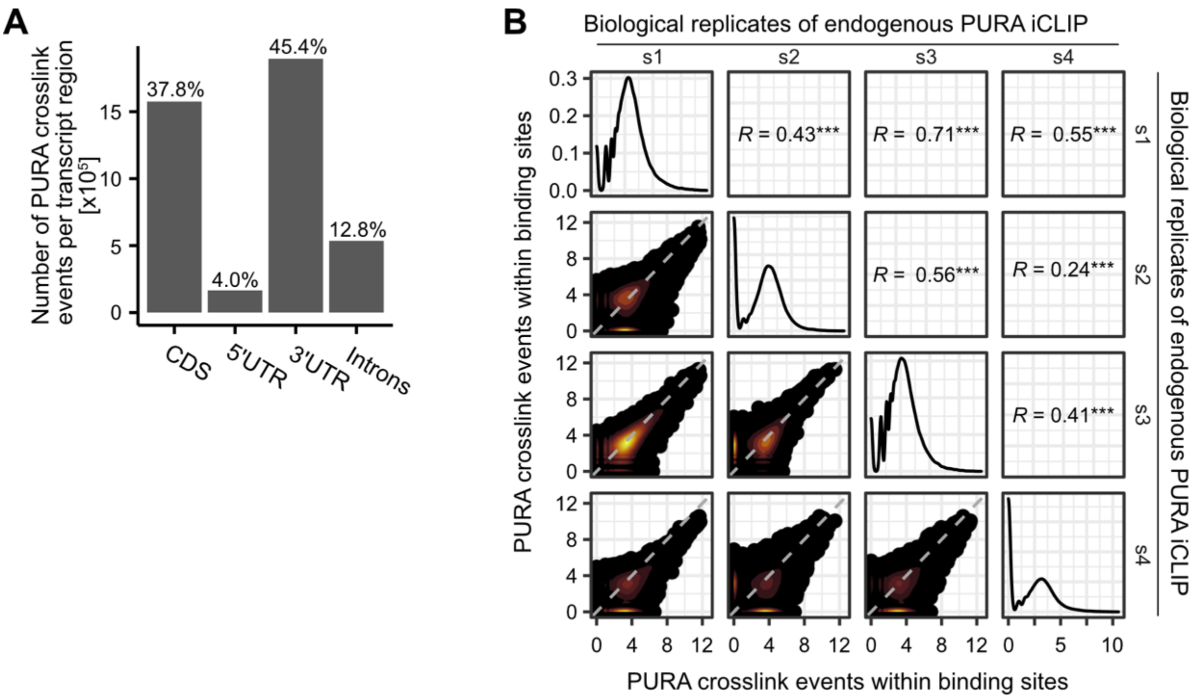

**Supplementary Figure S2: iCLIP shows global binding of endogenous PURA to RNAs. (A)** Distribution of PURA crosslink events in different transcript regions. Bars show the number of PURA crosslink events in the respective region, percentages depict the proportion from total crosslink events. **(B)** Correlation of PURA crosslink events per binding site between biological replicates is shown. Each plot compares two samples and each dot depicts a binding site. Pearson correlation coefficients (*R*) and associated *P* values are shown. ***, *P* value < 0.001.

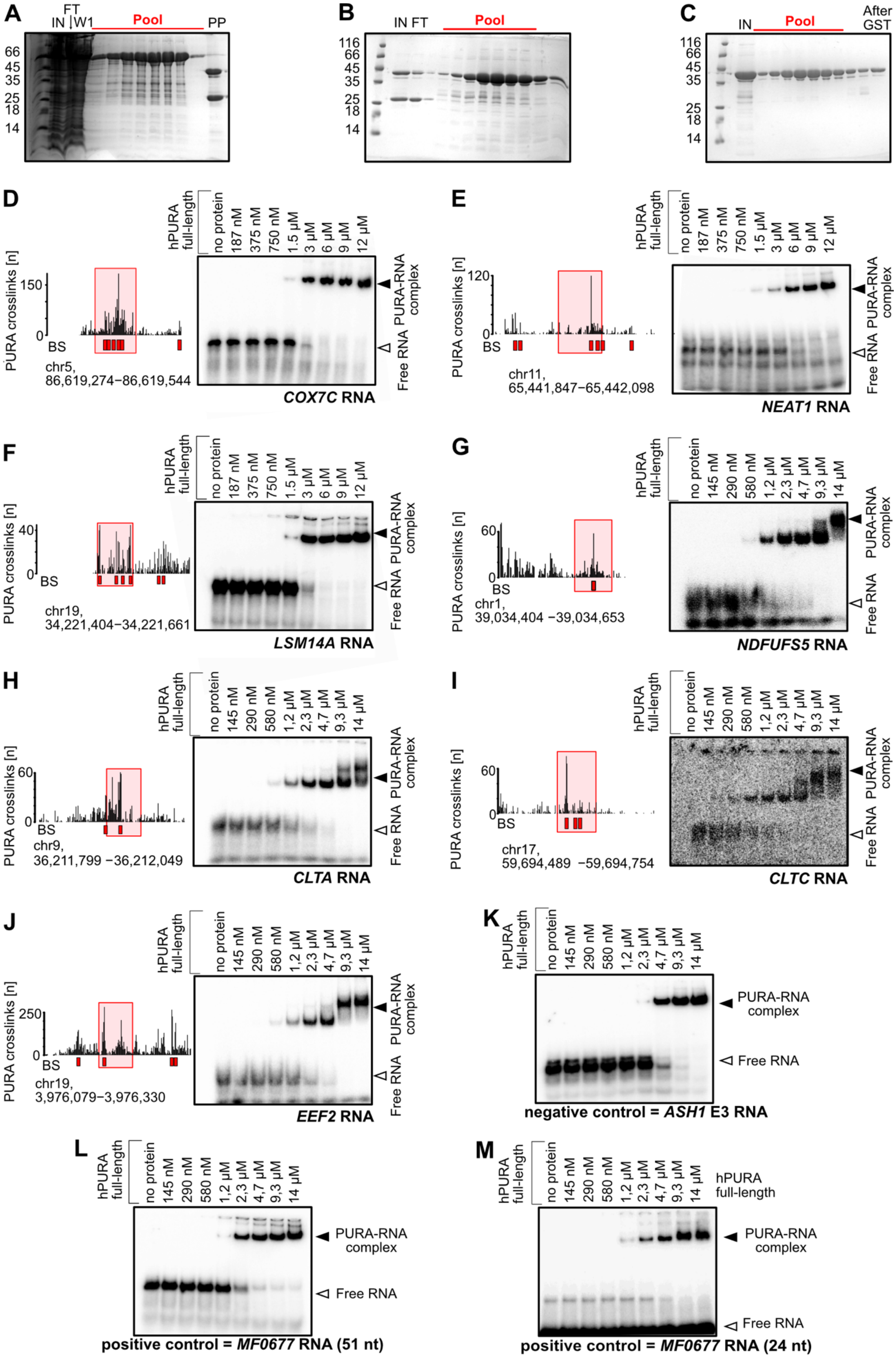

A
B
C
IN
FT
W1
Pool
PP
After GST
116
66
45
35
25
18
14
D
E
F
G
H
I
J
K
L
M
hPURA full-length
no protein
187 nM
375 nM
750 nM
1.5 µM
3 µM
6 µM
9 µM
12 µM
145 nM
290 nM
580 nM
1,2 µM
2,3 µM
4,7 µM
9,3 µM
14 µM
PURA crosslinks [n]
BS
PURA-RNA complex
Free RNA
chr5, 86,619,274−86,619,544
COX7C RNA
chr11, 65,441,847−65,442,098
NEAT1 RNA
chr19, 34,221,404−34,221,661
LSM14A RNA
chr1, 39,034,404 −39,034,653
NDFUFS5 RNA
chr9, 36,211,799 −36,212,049
CLTA RNA
chr17, 59,694,489 −59,694,754
CLTC RNA
chr19, 3,976,079−3,976,330
EEF2 RNA
negative control = ASH1 E3 RNA
positive control = MF0677 RNA (51 nt)
positive control = MF0677 RNA (24 nt)

**Supplementary Figure S3: EMSAs with iCLIP-identified target RNAs of PURA. (A-C)** Eluted fractions of subsequent purification steps, i.e., GST-TRAP column (A), heparin column (B), and size exclusion column (C), loaded on an SDS-PAGE. Marked is the pool used for the next step. The final concentration after size exclusion was 2.1 mg/ml. IN, input; FT, flow-through; W1, first wash step; PP, after precision protease cleavage; After GST, after further column purification with gst column. **(D-J)** Electrophoretic mobility shift assays (EMSA) show direct binding of PURA to target RNAs identified from PURA iCLIP experiments. Recombinantly expressed full-length human PURA was incubated with increasing concentrations (as indicated above) of radiolabeled RNA fragments (42-81nt) harboring PURA binding sites from the *COX7C* (D), *NEAT1* (E), *LSM14A* (F), *NDFUFS5* (G), *CLTA* (H), *CLTC* (I), and *EEF2* (J) RNAs. The bands corresponding to free RNA and formed PURA-RNA complexes are marked by white and black arrowheads, respectively. Genome browser views (left) show iCLIP signal for endogenous PURA (HeLa cells) in the region of the EMSA probe (red box). PURA binding sites are indicated below each track (BS). **(K-M)** Control EMSAs with *ASH1* RNA from yeast as unspecific binding partner (K) as well as the positive control *MF0677* RNA as a 51-mer (L) and 24-mer (M) as published in (5). All experiments were performed in triplicates and one representative EMSA gel is shown.

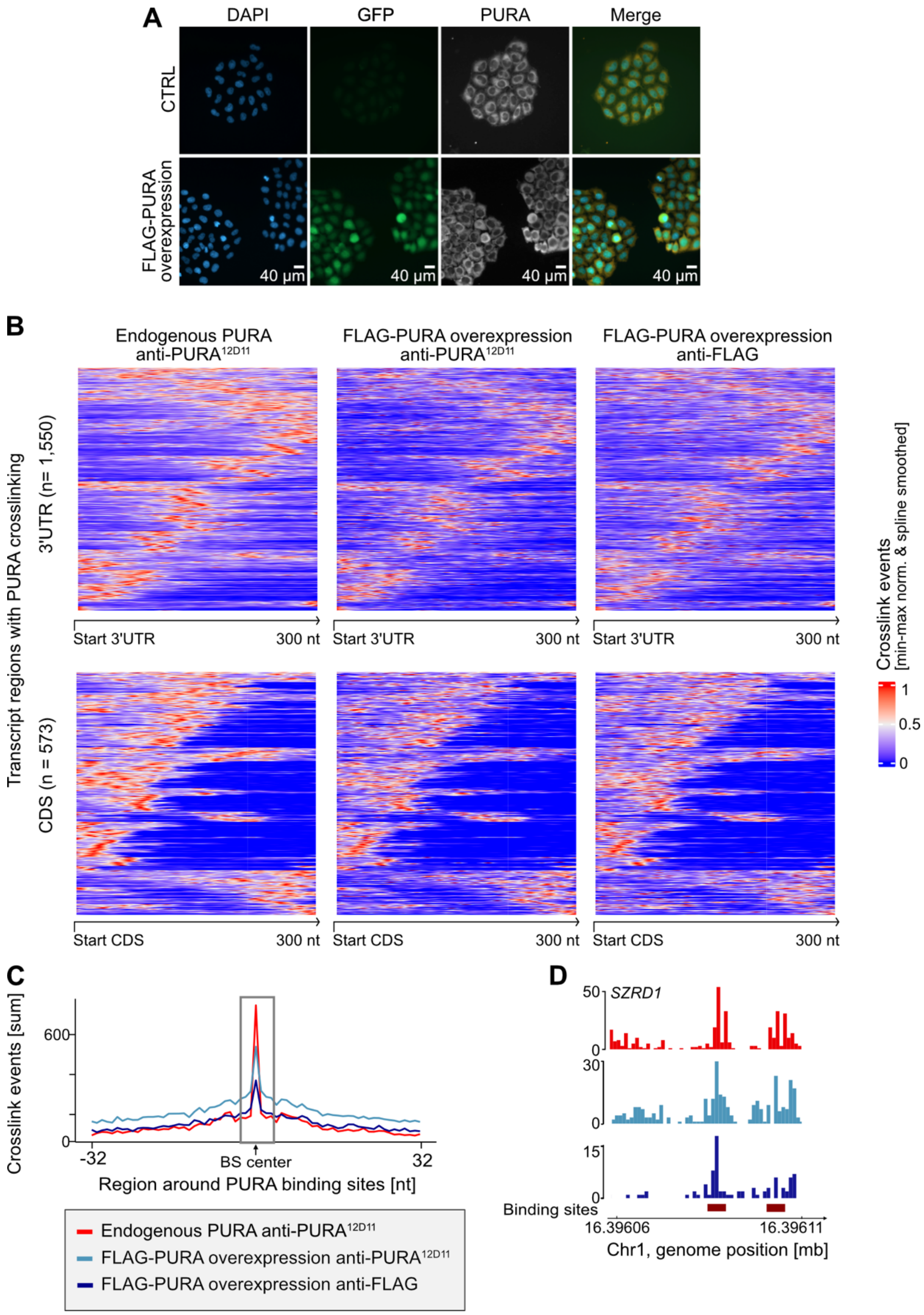

**Supplementary Figure S4: Comparison of PURA crosslink patterns from complementary iCLIP experiments validates specificity of the anti-PURA$^{12D11}$ antibody.** The PURA crosslink patterns from FLAG-PURA-overexpressing cells are comparable to the crosslink patterns from endogenous PURA, but contain a higher background signal. **(A)** HeLa cells with endogenous PURA levels (CTRL) and FLAG-PURA-P2A-GFP-overexpressing cells were stained for DAPI (blue), GFP (green) and

PURA (anti-PURA$^{12D11}$, 555 nm depicted in white). In the merge, PURA is shown in orange for better visualization. Scale bars, 40 µm. **(B)** PURA crosslink patterns in the first 300 nt of 3'UTRs (top) or CDS (bottom) with intermediate PURA crosslink coverage ($10^2$-$10^6$ crosslink events per window, 3'UTR: n = 1,550, CDS: n = 573). Each line depicts one 3`UTR or CDS. The crosslink intensity is given by the color scale. Crosslinks are min-max-normalized and spline-smoothed. While the peak regions (red) are similar in the different CLIP experiments, more background signal can be observed in the PURA overexpression experiments around these peak regions. **(C)** Metaprofile in 65-nt window around PURA binding sites (grey box) shows the sum of crosslink events for 1,000 randomly selected binding sites (red – endogenous PURA, anti-PURA$^{12D11}$; lightblue – FLAG-PURA overexpression, anti-PURA$^{12D11}$; darkblue – FLAG-PURA overexpression, anti-FLAG). Crosslink events are min-max-normalized and spline-smoothed. Background signal around binding sites is higher in PURA overexpression experiments. **(D)** Exemplary PURA crosslink events on the *SZRD1* RNA (color coding as in (C), boxes in darkred indicate binding sites defined for endogenous PURA iCLIP).

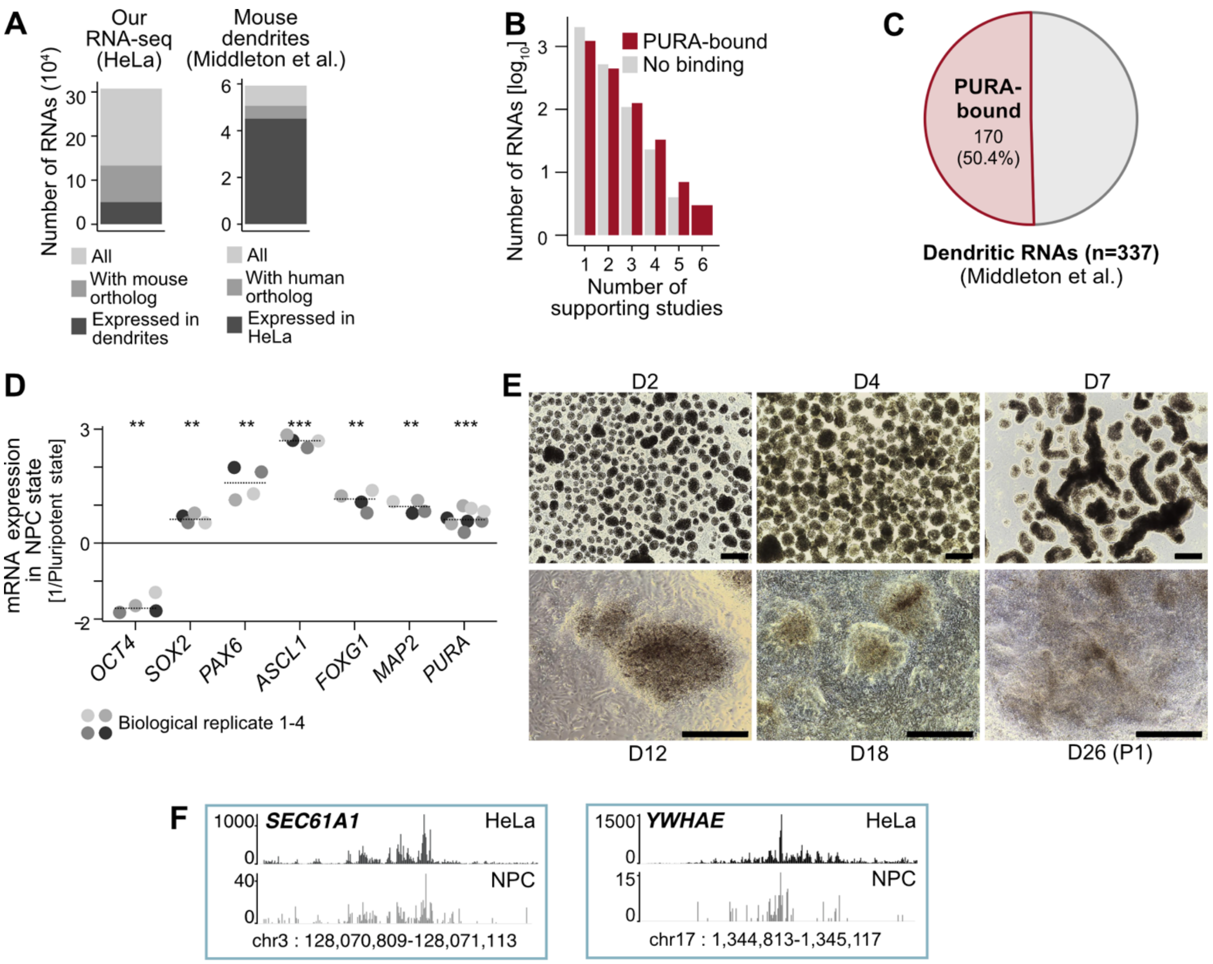

**Supplementary Figure S5: Validation of results in neural precursor cells (NPCs).** **(A)** To overlap PURA-bound RNAs from human HeLa cells with dendritically localized RNAs from mouse (6), all genes present in either dataset ("All") were filtered for having a 1-to-1 ortholog in the other species and being expressed in the other dataset. Shown is the number of RNAs that met these criteria both from the perspective of our HeLa cell data (left) and the mouse dendritic RNAs from (6) (right). **(B)** Overlap of PURA-bound RNAs with dendritically localized RNAs from (6) stratified by minimum number of supporting studies. **(C)** Pie chart shows overlap of PURA-bound transcripts with dendritically localized RNAs identified in at least 3 out of 6 independent studies in (6) (n = 337). **(D)** Expression of RNAs that mark transition from human induced pluripotent stem cells (hiPSCs) to NPCs measured by qPCR. Expression of all RNAs is normalized to *GAPDH*, and expression in NPCs is given relative to expression in hiPSCs. Dots show four biological replicates and dotted line mean of those. *PURA* expression was measured with two different primer sets. **, $P$ value < 0.01, ***, $P$ value < 0.001, unpaired, two-sided Student's $t$-test. **(E)** Different time points during NPC differentiation from hiPSCs to NPCs. Morphological changes through differentiation checkpoints were observed as expected. Time points of image acquisition are shown in days (scale bars, 500 µm). **(F)** Genome browser views illustrate exemplary binding sites on the *SEC61A1* (left) and *YWHAE* (right) transcripts in the HeLa (darkgrey) and NPC (lightgrey) iCLIP data.

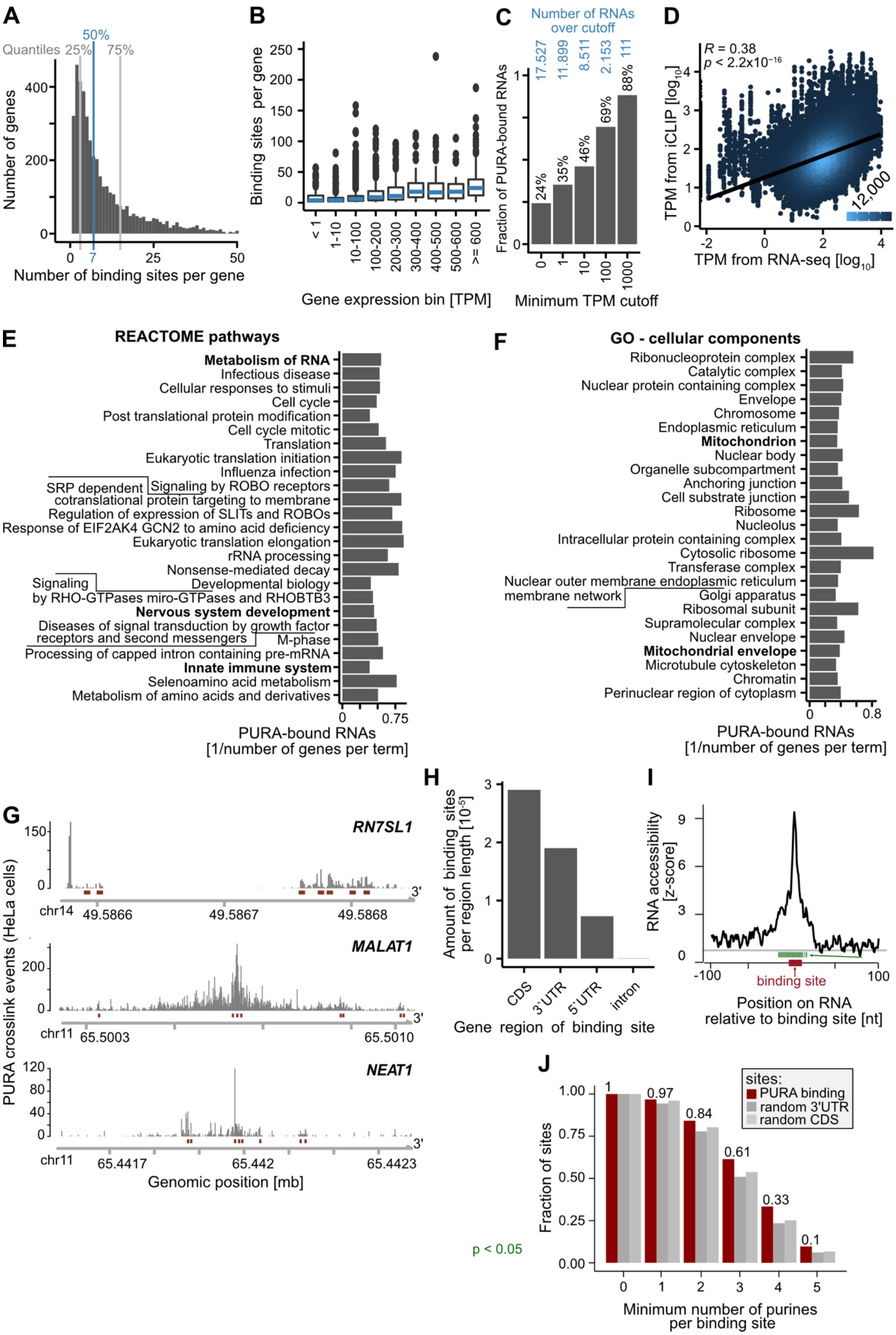
A
Quantiles 25%
50%
75%
Number of genes
400
200
0
0
7
25
50
Number of binding sites per gene
B
Binding sites per gene
250
200
150
100
50
0
< 1
1-10
10-100
100-200
200-300
300-400
400-500
500-600
>= 600
Gene expression bin [TPM]
C
Number of RNAs over cutoff
17.527
11.899
8.511
2.153
111
Fraction of PURA-bound RNAs
1
0
24%
35%
46%
69%
88%
0
1
10
100
1000
Minimum TPM cutoff
D
R = 0.38
p < 2.2x10^-16
TPM from iCLIP [log10]
TPM from RNA-seq [log10]
12,000
E
REACTOME pathways
Metabolism of RNA
Infectious disease
Cellular responses to stimuli
Cell cycle
Post translational protein modification
Cell cycle mitotic
Translation
Eukaryotic translation initiation
Influenza infection
Signaling by ROBO receptors
SRP dependent cotranslational protein targeting to membrane
Regulation of expression of SLITs and ROBOs
Response of EIF2AK4 GCN2 to amino acid deficiency
Eukaryotic translation elongation
rRNA processing
Nonsense-mediated decay
Developmental biology
Signaling by RHO-GTPases miro-GTPases and RHOBTB3
Nervous system development
Diseases of signal transduction by growth factor receptors and second messengers
M-phase
Processing of capped intron containing pre-mRNA
Innate immune system
Selenoamino acid metabolism
Metabolism of amino acids and derivatives
0
0.75
PURA-bound RNAs
[1/number of genes per term]
F
GO - cellular components
Ribonucleoprotein complex
Catalytic complex
Nuclear protein containing complex
Envelope
Chromosome
Endoplasmic reticulum
Mitochondrion
Nuclear body
Organelle subcompartment
Anchoring junction
Cell substrate junction
Ribosome
Nucleolus
Intracellular protein containing complex
Cytosolic ribosome
Transferase complex
Nuclear outer membrane endoplasmic reticulum membrane network
Golgi apparatus
Ribosomal subunit
Supramolecular complex
Nuclear envelope
Mitochondrial envelope
Microtubule cytoskeleton
Chromatin
Perinuclear region of cytoplasm
0
0.8
PURA-bound RNAs
[1/number of genes per term]
G
PURA crosslink events (HeLa cells)
RN7SL1
150
0
3'
chr14
49.5866
49.5867
49.5868
MALAT1
200
0
3'
chr11
65.5003
65.5010
NEAT1
120
80
40
0
3'
chr11
65.4417
65.442
65.4423
Genomic position [mb]
H
Amount of binding sites per region length [10^-5]
3
2
1
0
CDS
3'UTR
5'UTR
intron
Gene region of binding site
I
RNA accessibility [z-score]
9
6
3
0
-100
100
binding site
Position on RNA relative to binding site [nt]
J
sites:
PURA binding
random 3'UTR
random CDS
Fraction of sites
1.00
0.75
0.50
0.25
0.00
1
0.97
0.84
0.61
0.33
0.1
p < 0.05
0
1
2
3
4
5
Minimum number of purines per binding site

**Supplementary Figure S6: Accessibility and sequence composition at PURA binding sites. (A)** Distribution of PURA binding sites on PURA-bound genes. Histogram shows number of binding sites per gene against number of genes. Lines annotate the median (blue), lower quartile and upper quartile (grey). **(B)** The influence of the gene expression (estimated as transcripts per million [TPM]) on the number of binding sites per gene. Genes are stratified into bins by their TPM displayed on the x-axis. The number of binding sites per gene is displayed in a boxplot for all genes in each bin. **(C)** Fraction of PURA bound transcripts increases with the expression levels of the RNAs. Bars show the fraction of PURA bound transcripts against cutoff of expression of RNAs measured as transcripts per million (TPM, based on RNA-seq). Total number of RNAs above cutoff is given above the bar. For instance, at a TPM cutoff > 10, 3,951 out of 8,511 transcripts (46%) are bound by PURA. **(D)** TPM values can be reliably estimated from total iCLIP crosslink events or RNA-seq data. Shown is a scatterplot of TPM values calculated from RNAseq data (x-axis) versus iCLIP data (y-axis). Pearson correlation coefficient and associated *P* value are given. **(E, F)** Top 25 enriched REACTOME pathways (E) and Gene Ontology cellular components (F) for PURA bound transcripts. Bars show the ratio of target RNAs to the total number of genes per term. All *P* values are below 0.0001. **(G)** Genome browser view shows the pileup of crosslink events of endogenous PURA from HeLa cells at binding sites (red) in the long non-coding RNAs *RN7SL1*, *MALAT1* and *NEAT1*. **(H)** PURA binding sites in different transcript regions of protein-coding genes relative to the length of the regions in all expressed transcripts in percent. **(I)** Prediction of accessibility of RNA around PURA binding sites shows an increased propensity of the bindings sites to be single-stranded (i.e., higher RNA accessibility). A z-score of the predicted accessibility per position over a background of random regions (y-axis) is shown along a 201-nt window around the binding sites (x-axis). RNA accessibility was predicted by RNAplfold (7) in a window of 501 nt around the PURA binding sites from which the center 201 nt are displayed. Green lines on x-axis indicate positions with adjusted *P* value < 0.05. **(J)** PURA binding sites contain accumulations of purines. Shown is the fraction of binding sites (red bar and number given above) containing at least 0-5 purines (given on the x-axis) in comparison to random sites in the 3'UTR (darkgrey) or CDS (lightgrey).

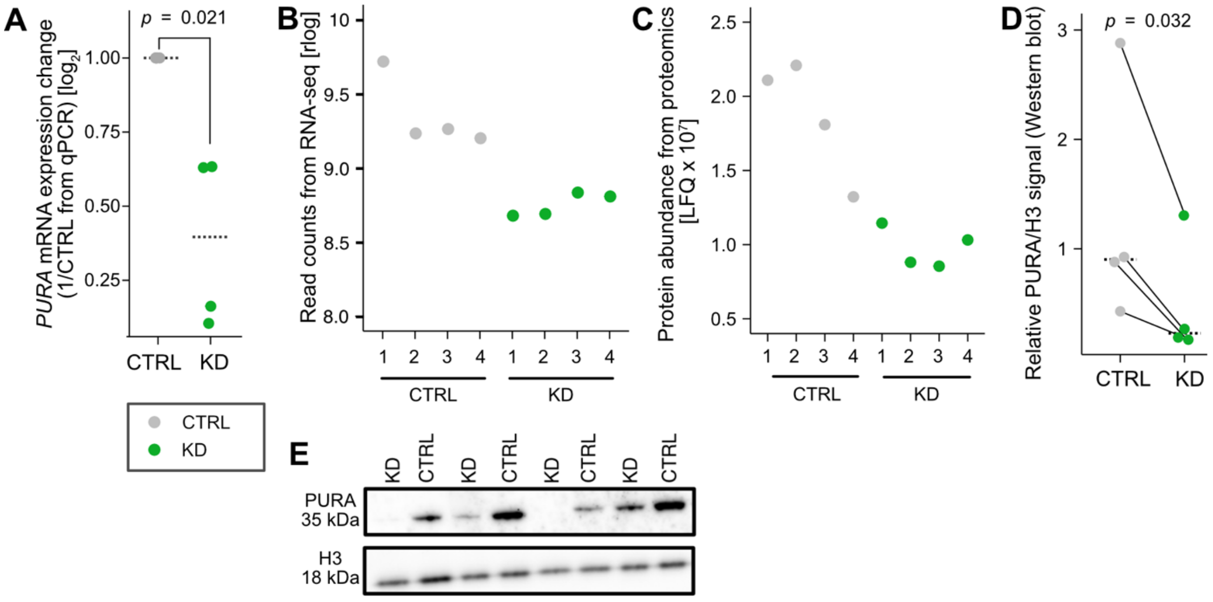

**Supplementary Figure S7: Validation of siRNA-mediated *PURA* knockdown.** **(A)** Relative quantification of *PURA* qPCR signal from *PURA* KD and CTRL HeLa cells (n = 4 biological replicates), *P* value from unpaired two-sided Student's *t*-test. **(B)** Normalized PURA read counts (rlog, regularized log transformation by DESeq2 (8)) in *PURA* KD (green) and CTRL (grey) HeLa cell samples from RNAseq experiment. **(C)** PURA protein abundances in *PURA* KD (green) and CTRL (grey) HeLa cell samples from shotgun proteomics experiment. Abundances are normalized to LFQ. (**D, E)** Western blot experiments (E) and quantification (D) on samples to validate *PURA* KD for shot-gun proteomics, *P* value from one-sided paired Student's *t*-test.

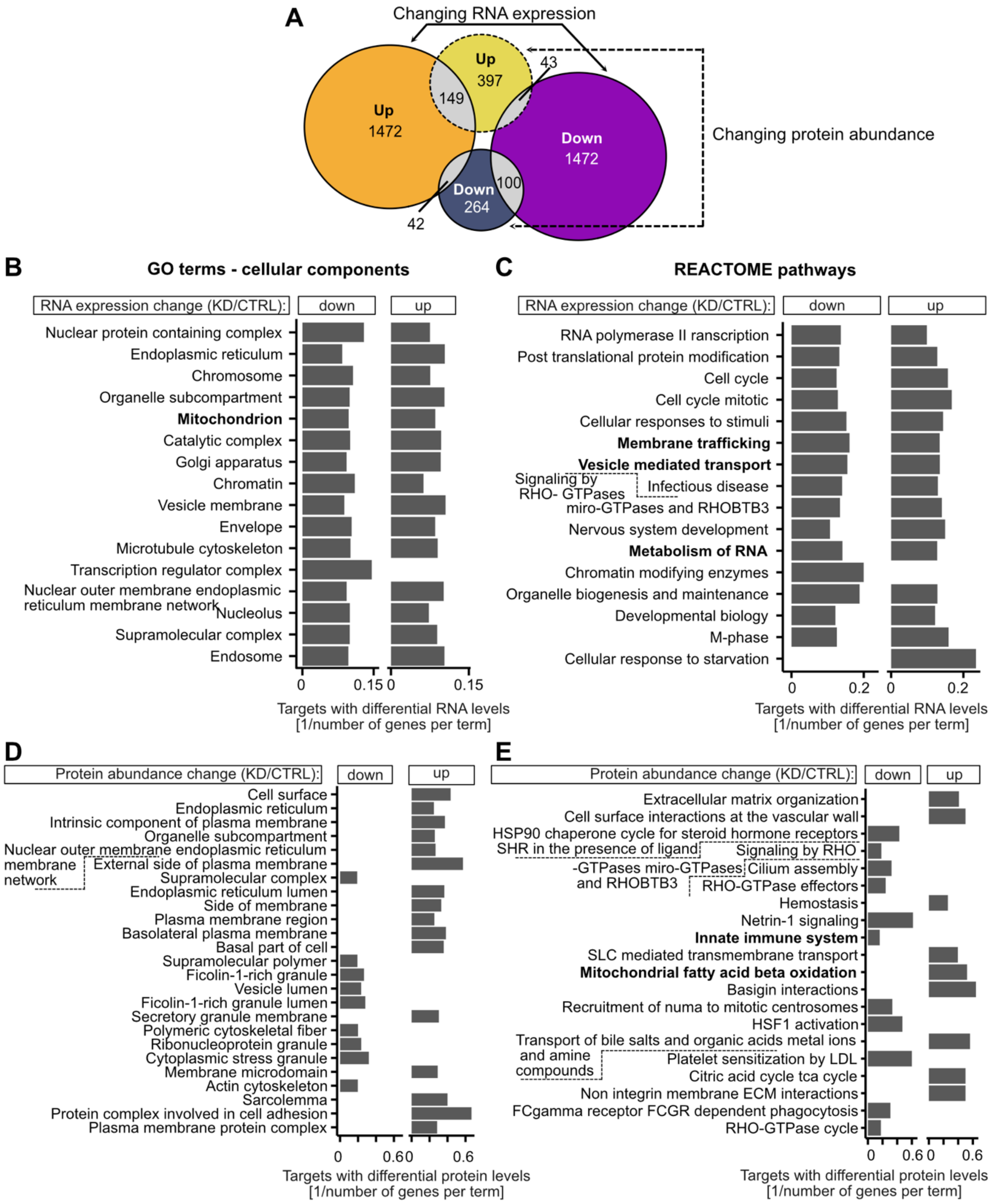

**Supplementary Figure S8: Functional enrichment analyses of differentially expressed RNAs and proteins upon *PURA* KD.** **(A)** Overlap of genes that are significantly up- or downregulated at the RNA level (solid line, up – orange, down – purple, FDR < 0.01) and protein level (dashed line, up - yellow, down – darkblue, FDR < 0.05) in *PURA* KD/CTRL RNA-seq and proteomics experiments, respectively. **(B-E)** Functional enrichment analysis of up- and downregulated genes/proteins that significantly change in *PURA* KD compared to CTRL condition in the RNA-seq (B, C; FDR < 0.01) and proteomics (D, E; FDR < 0.05) experiments. Top 20 Gene Ontology

(GO) terms for cellular component (B, D) and top 20 enriched REACTOME pathways (C, E) are shown. Bars show the ratio of target RNAs with significant differential regulation to the total number of genes per term.

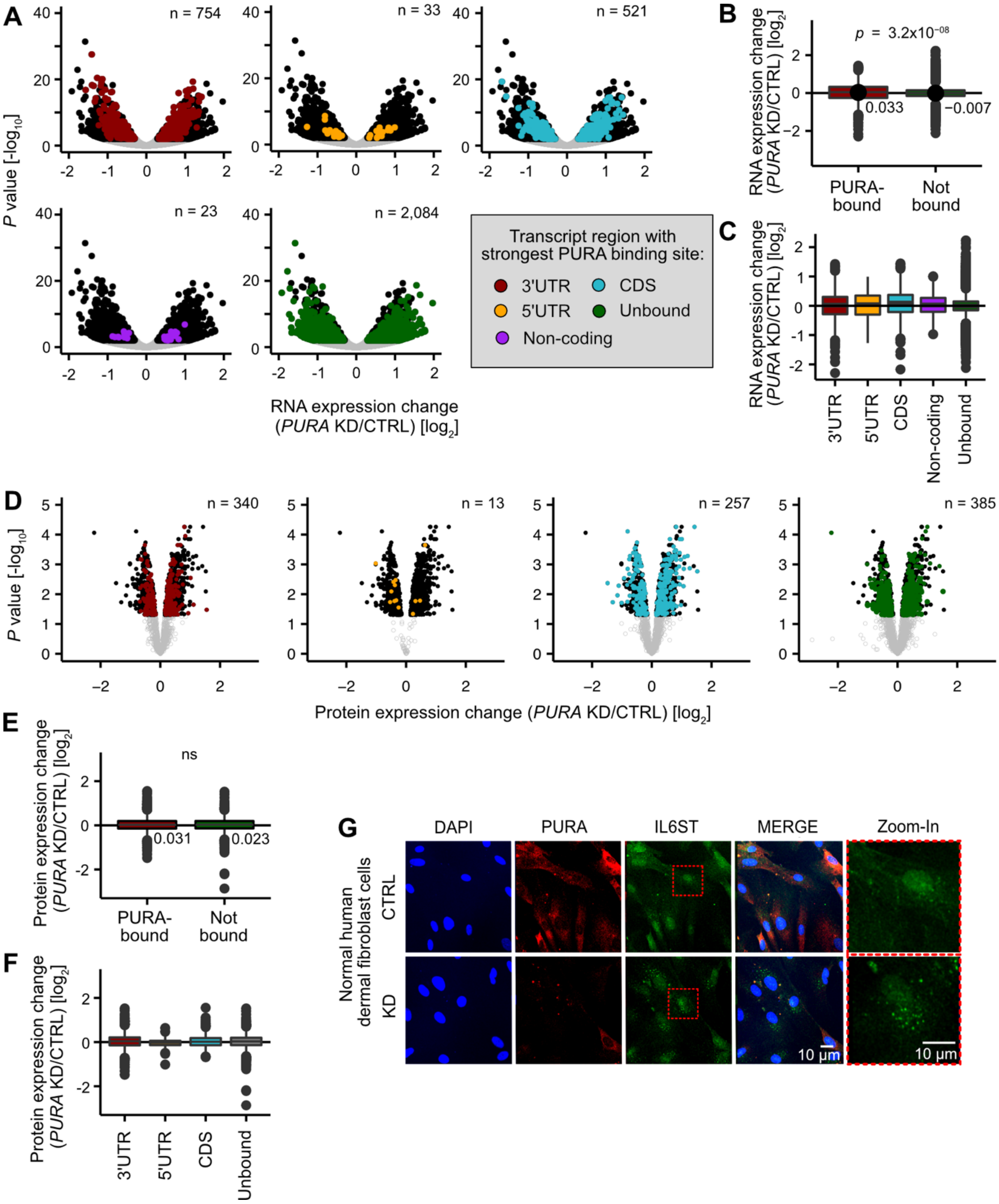

**Supplementary Figure S9: The changes in RNA and protein levels upon *PURA* KD are not linked to PURA binding in a particular transcript region.** **(A)** Volcano plots showing differentially expressed genes in *PURA* knockdown (KD) *versus* control (CTRL) HeLa cells (black: 3,415 significantly changing genes with FDR < 0.01) colored by transcript region with the strongest PURA binding site (3'UTR, red; 5'UTR, orange; CDS, turquoise; non-coding, violet; unbound RNAs, green). **(B)** Distribution of expression changes of PURA-bound (red) and unbound (green) RNAs. Mean $\log_2$ fold changes (*PURA* KD/CTRL) are given. *P* value = $4.9\text{x}10^{-06}$, unpaired two-sided Student's *t*-test. **(C)** Distribution of expression changes as in (B) with PURA-bound

RNAs stratified by transcript region with strongest PURA binding site. Color coding as in (A). **(D)** Volcano plots showing differentially expressed proteins in *PURA* knockdown (KD) *versus* control (CTRL) HeLa cells (black: 995 significantly changing proteins with FDR < 0.05) colored by transcript region with the strongest PURA binding site. Color coding as in (A). **(E)** Distribution of expression changes of PURA-bound (red) and unbound (green) proteins. Mean $\log_2$ fold changes (*PURA* KD/CTRL) are given. *P* value is non-significant (ns), unpaired two-sided Student's *t*-test. **(F)** Distribution of expression changes as in (B) with PURA-bound proteins stratified by transcript region with strongest PURA binding site. Color coding as in (A). **(G)** *PURA* KD leads to increased IL6ST expression and intracellular granule formation in normal human dermal fibroblast (NHDF) cells. Immunofluorescence staining of PURA (anti-PURA$^{12D11}$, red) and IL6ST (green), as well as DAPI staining (blue) in CTRL and *PURA* KD NHDF cells. Scale bars, 10 µm.

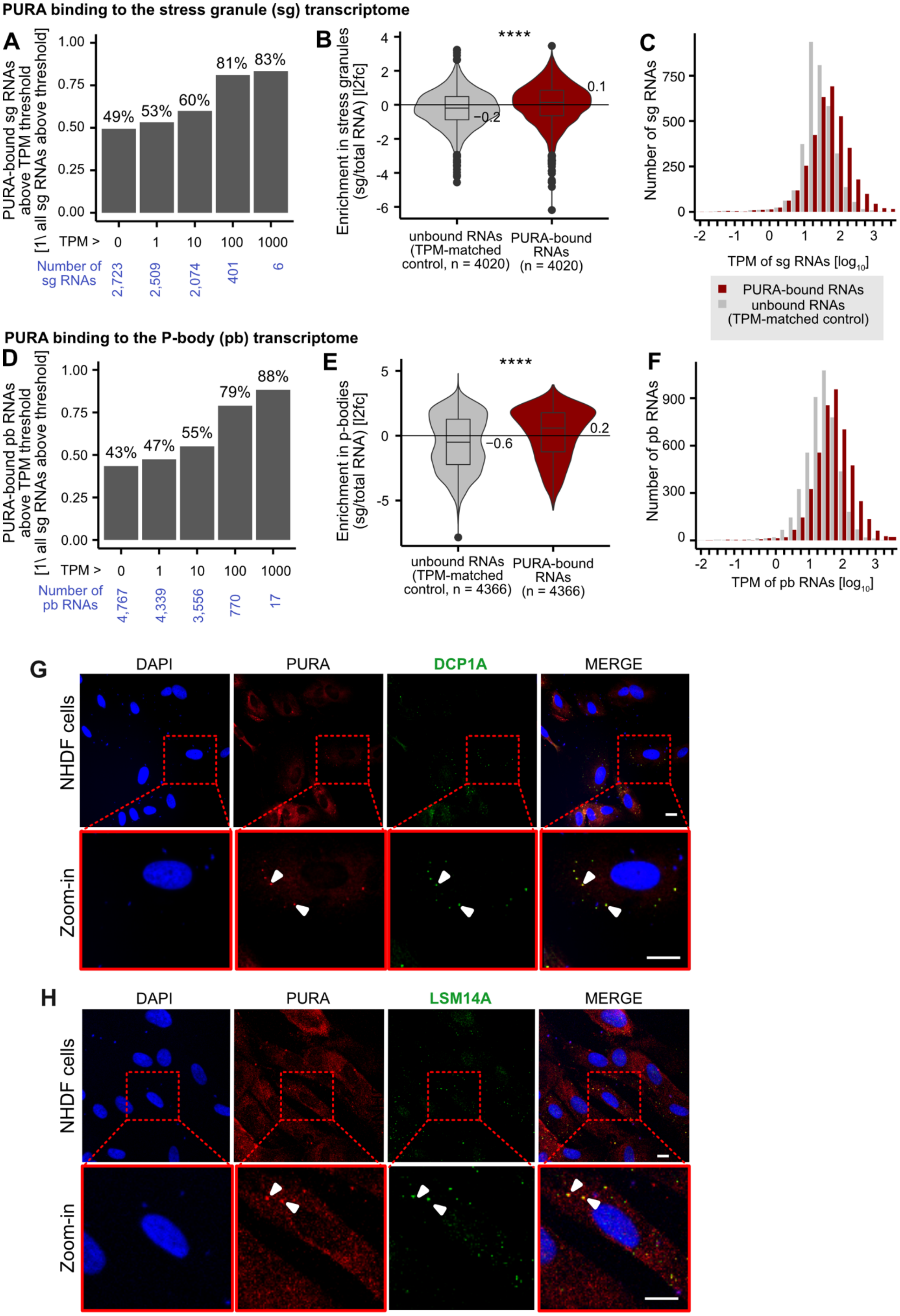
PURA binding to the stress granule (sg) transcriptome
A
PURA-bound sg RNAs above TPM threshold [1\ all sg RNAs above threshold]
49%
53%
60%
81%
83%
TPM >
0
1
10
100
1000
Number of sg RNAs
2,723
2,509
2,074
401
6
B
Enrichment in stress granules (sg/total RNA) [l2fc]
****
−0.2
0.1
unbound RNAs (TPM-matched control, n = 4020)
PURA-bound RNAs (n = 4020)
C
Number of sg RNAs
TPM of sg RNAs [log10]
PURA-bound RNAs
unbound RNAs (TPM-matched control)
PURA binding to the P-body (pb) transcriptome
D
PURA-bound pb RNAs above TPM threshold [1\ all sg RNAs above threshold]
43%
47%
55%
79%
88%
TPM >
0
1
10
100
1000
Number of pb RNAs
4,767
4,339
3,556
770
17
E
Enrichment in p-bodies (sg/total RNA) [l2fc]
****
−0.6
0.2
unbound RNAs (TPM-matched control, n = 4366)
PURA-bound RNAs (n = 4366)
F
Number of pb RNAs
TPM of pb RNAs [log10]
G
DAPI
PURA
DCP1A
MERGE
NHDF cells
Zoom-in
H
DAPI
PURA
LSM14A
MERGE
NHDF cells
Zoom-in

**Supplementary Figure S10: PURA-bound RNAs are likely to be enriched in the stress granule and p-body transcriptome and PURA localizes in P-bodies in normal human dermal fibroblast (NHDF) cells.** **(A-F)** Comparison of PURA-bound RNAs to previously published stress granule (sg) and P-body (pb) transcriptomes, defined as RNAs significantly enriched in the given granule (FDR < 0.01, log2foldChange > 0) as calculated by (9) (sg) and (10) (pb). We considered only RNAs which were expressed in our PURA iCLIP experiment from HeLa cells as well as the U-2 OS cells from (9) (n = 2,723) or the HEK293 cells from (10) (n = 4,767). (A, D) Intersection of PURA-bound RNAs with the sg (A) or pb (D) transcriptome in relation to their RNA expression level in HeLa cells. The minimum RNA expression is given as TPM on the x-axis. Number of RNAs is given below (blue). The y-axis and numbers over the bars specify the ratio of PURA-bound to non-bound RNAs. (B, E) Distributions of enrichment values for PURA-bound RNAs (red) in sg (B) and pb (E) taken from by (9) (sg) and (10) (pb) (mean enrichment is given next to violins). In a comparison to a control set of RNAs not bound by PURA (grey), PURA-bound RNAs are significantly more enriched in both sg and pb (*P* value < 0.0001, unpaired two-sided Wilcoxon Rank-sum test). To account for the influence of the RNA expression levels on the binding detection, a TPM-adjusted control set was used. (C, F) TPM distribution of PURA-bound RNAs and control set after TPM adjustment of the control set for sg (B) and pb (E). **(G, H)** Colocalization confirms P-body association of PURA in NHDF cells as seen in HeLa cells (**Figure 5B, C**). Immunofluorescence staining of P-body marker proteins (green) DCP1A (G) and LSM14A (H) together with PURA (anti-PURA$^{12D11}$, red) and DAPI (blue). Arrowheads indicate examples of granules with overlapping staining. Scale bars, 10 µm.

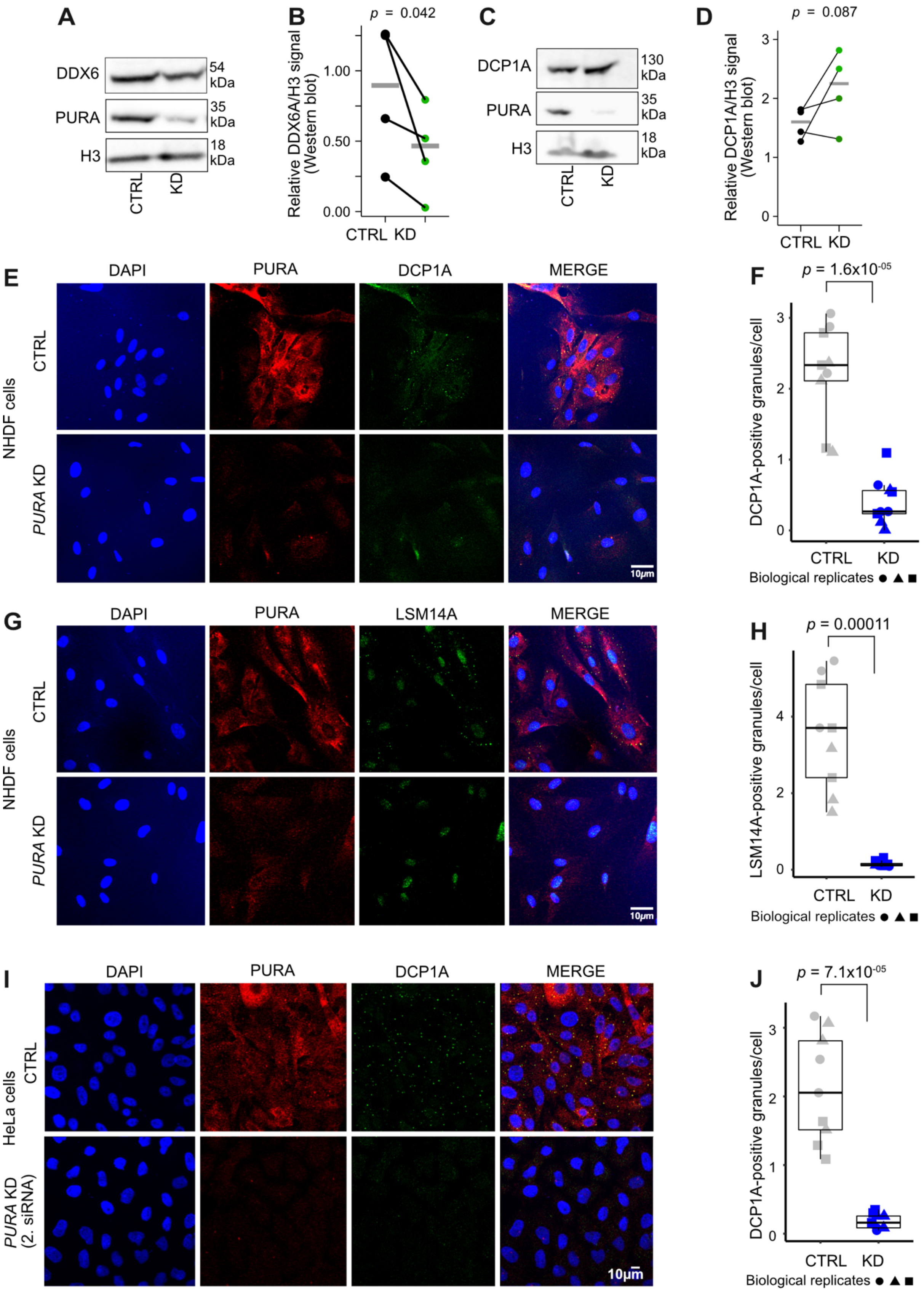

**Supplementary Figure S11: *PURA* KD leads to a loss of P-bodies in NHDF cells as seen in HeLa cells (Figure 6C-H).** **(A, B)** DDX6 levels are significantly downregulated upon *PURA* KD. Representative Western blot (A) and quantification (B) (n = 4) are shown for DDX6 in CTRL and *PURA* KD HeLa cells (n = 4). *P* value from unpaired one-sided Student's *t*-test. **(C, D)** DCP1A levels do not significantly change upon *PURA* KD. Representative Western blot (C) and quantification (D) are shown for DCP1A in CTRL and *PURA* KD HeLa cells (n = 4). *P* value from unpaired one-sided Student's *t*-test. **(E, G)** Immunofluorescence staining in NHDF cells of PURA (anti-PURA$^{12D11}$, red) together with P-body marker proteins (measured in 555 nm channel, depicted in green) DCP1A (E) and LSM14A (G) as well as DAPI (blue) as a nuclear stain in CTRL (top) and *PURA* KD (bottom). Scale bars, 10 µm. **(F, H)** ImageJ analysis of DCP1A-positive (F) and LSM14A-positive (H) granules per cell on approximately 400 cells per replicate (n = 3 [DCP1A] and 3 [LSM14A] replicates) comparing CTRL and *PURA* KD conditions. Granules were defined as described in Methods. *P* values from unpaired two-sided Student's *t*-test. **(I)** Validation of loss of P-bodies in HeLa cells upon *PURA* KD with an independent siRNA. Immunofluorescence staining of PURA (anti-PURA$^{12D11}$, red) together with P-body marker protein DCP1A (measured in 555 nm channel, depicted in green) as well as DAPI (blue) as a nuclear stain in CTRL (top) and *PURA* KD (bottom). Scale bars, 10 µm. **(J)** ImageJ analysis of DCP1A-positive granules form (I) as in (F).

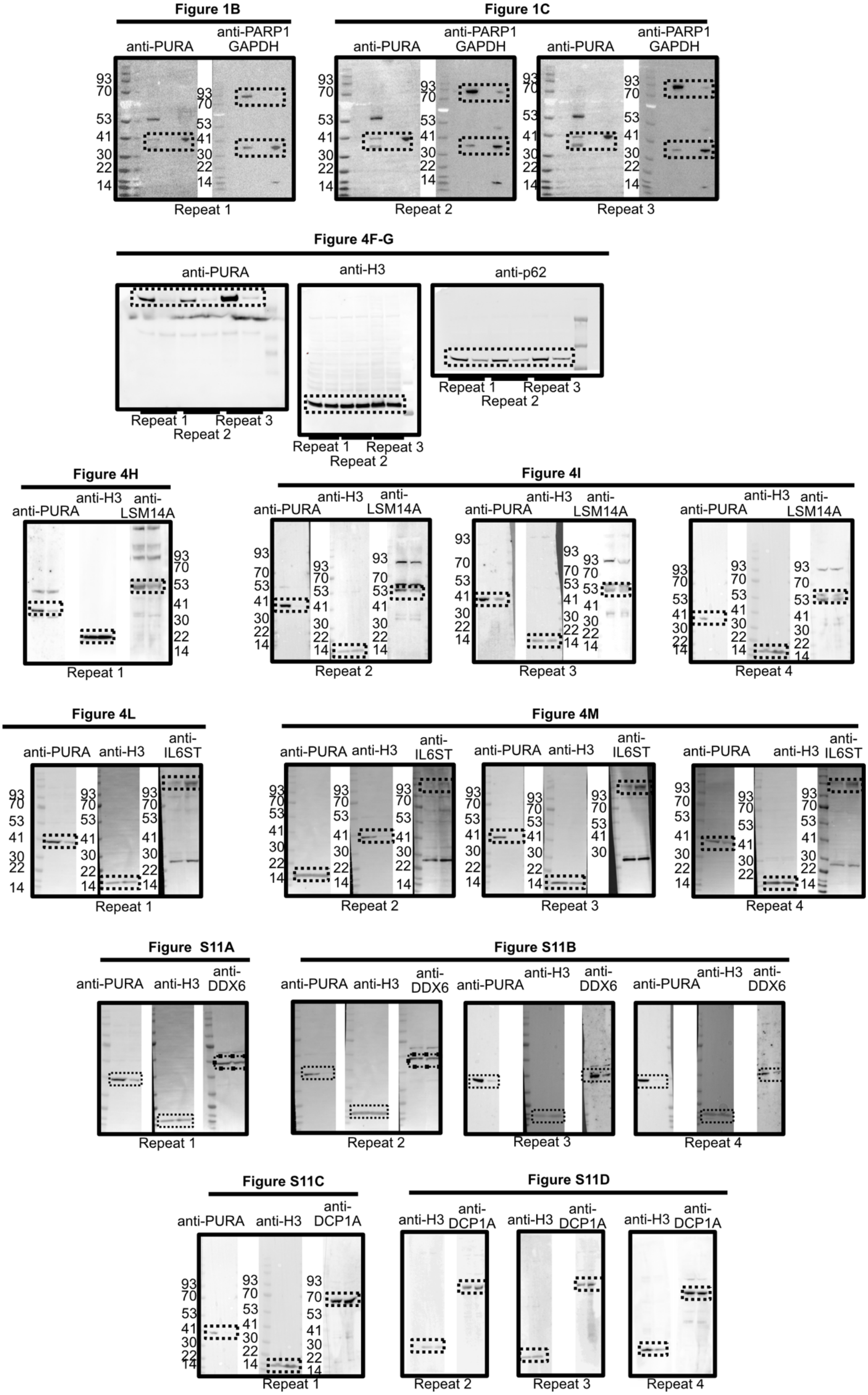
Figure 1B
anti-PURA
anti-PARP1 GAPDH
Repeat 1
Figure 1C
anti-PURA
anti-PARP1 GAPDH
Repeat 2
Repeat 3
93
70
53
41
30
22
14
Figure 4F-G
anti-PURA
anti-H3
anti-p62
Repeat 1
Repeat 2
Repeat 3
Figure 4H
anti-PURA
anti-H3
anti-LSM14A
Figure 4I
Repeat 4
Figure 4L
anti-IL6ST
Figure 4M
Figure S11A
anti-DDX6
Figure S11B
Figure S11C
anti-DCP1A
Figure S11D

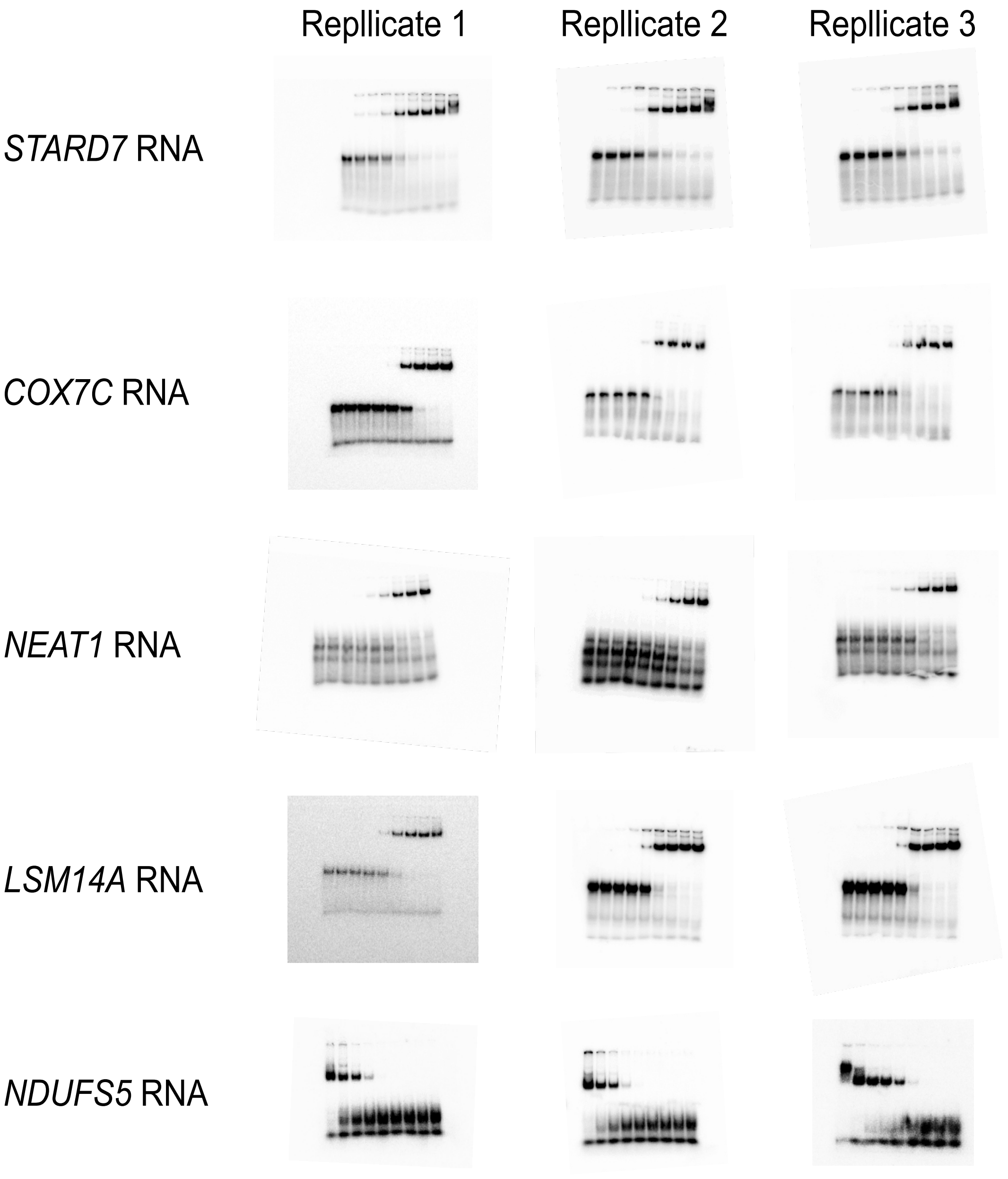
Repllicate 1
Repllicate 2
Repllicate 3
*STARD7* RNA
*COX7C* RNA
*NEAT1* RNA
*LSM14A* RNA
*NDUFS5* RNA

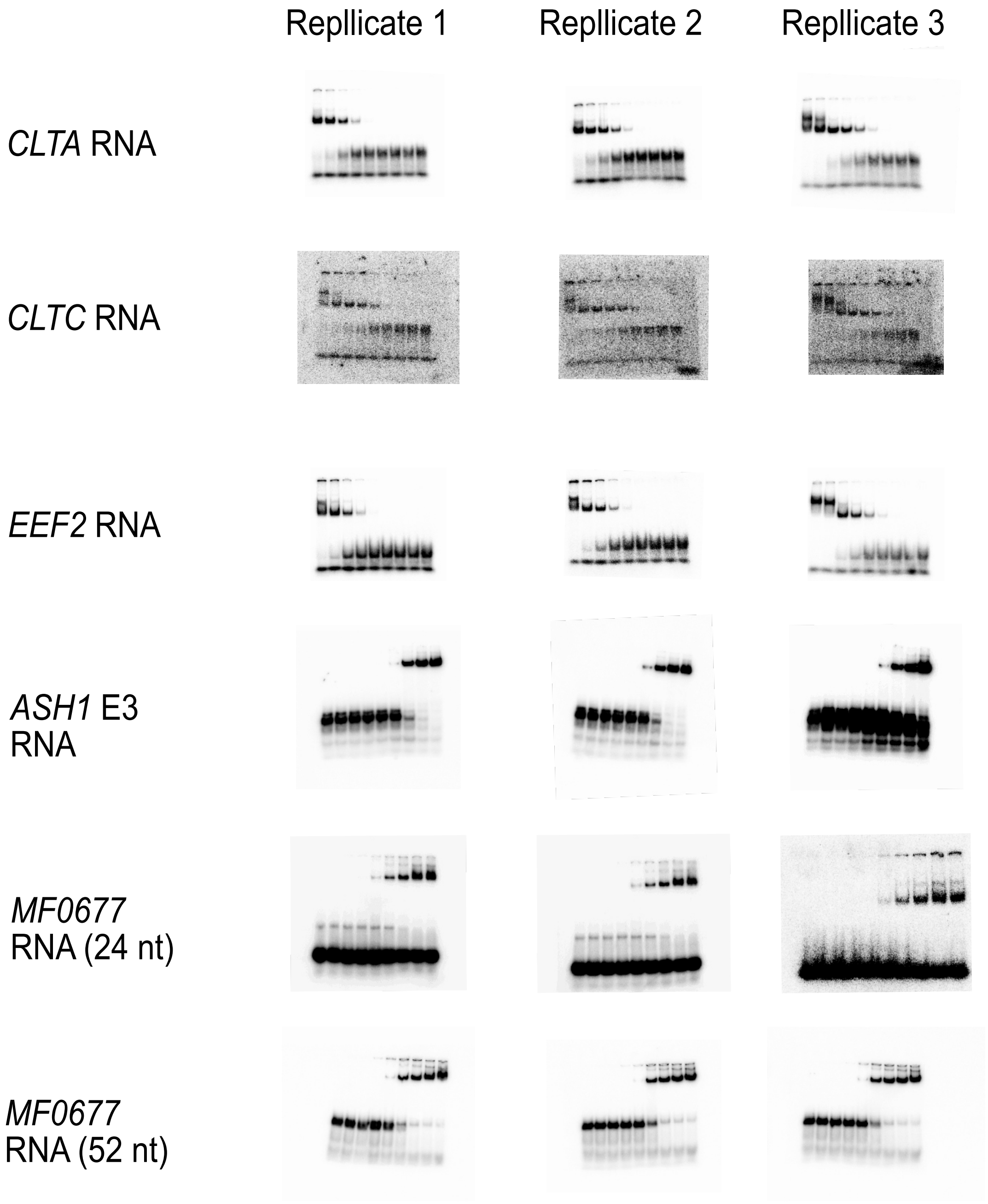

**Supplementary Figure S12: uncropped Western blots and EMSAs**

### Supplementary Material

#### Supplementary Material 1: Comparison of PURA crosslink patterns from complementary iCLIP experiments

This study contains four complementary iCLIP experiments for PURA performed at two different PURA levels (endogenous and FLAG-PURA overexpression), using two different antibodies (anti-PURA$^{12D11}$ and anti-FLAG) and two different cell lines (HeLa and NPC). Here, we describe the comparison between data sets and their implications in more detail.

<u>1) iCLIP experiments with overexpressed FLAG-PURA immunoprecipitated with anti-PURA$^{12D11}$ and anti-FLAG lead to comparable crosslink patterns, supporting the specificity of the anti-PURA$^{12D11}$ antibody.</u>

We performed two complementary iCLIP experiments with ectopic FLAG-PURA expression using a stable overexpressing HeLa cell line, resulting in an increase in PURA concentration in comparison to normal HeLa cells. We controlled the correct localization of the FLAG-PURA construct by immunofluorescence (IF) staining (**Supplementary Figure S4A**). In visual inspection, the obtained crosslink patterns look very similar as exemplarily shown in **Supplementary Figure S4D**. To compare the crosslink patterns in both experiments more broadly, we randomly selected 1,000 3'UTR and CDS windows and compared the PURA iCLIP signal therein after min-max normalization and spline smoothing. Indeed, the normalized crosslink patterns obtained from both iCLIP experiments were very similar as shown in **Supplementary Figure S4B**, supporting that our in-house anti-PURA$^{12D11}$ antibody is specific for PURA in immunoprecipitation experiments such as iCLIP.

<u>2) The increase of cellular PURA levels by FLAG-PURA overexpression is accompanied to a higher background signal</u>

The two iCLIP experiments with ectopic FLAG-PURA expression also allow us to compare the binding behavior of endogenous PURA to PURA overexpression. We found that the peaks in the crosslink signal of endogenous PURA generally matched those in the two overexpression experiments (**Supplementary Figure S4B**, red coloring), but were surrounded by a lower background signal (blue vs white coloring). This was further supported by a metaprofile of summed normalized crosslink events from the three experiments around each binding site (**Supplementary Figure S4C**): The crosslink profiles from all three experiments peaked in the center of the defined binding sites, indicating that overexpressed PURA also recognizes the binding sites that we obtained from the endogenous PURA iCLIP. At the same time, the relationship of signal in vs. around PURA binding sites is best for the endogenous PURA iCLIP. We hypothesize that the higher background signal in PURA overexpression arises due to unspecific RNA binding by the unphysiologically abundant ectopic PURA. For this reason, we used the endogenous PURA iCLIP for all further analyses.

3) PURA binds to similar binding sites in neuronal precursor cells

To test for PURA binding in a physiological cell system, we transferred our PURA binding analyses to neuronal precursor cells (NPCs). However, since PURA is only moderately expressed in these cells, a high amount of cell material had to be collected and we did not obtain a sufficient iCLIP signal for endogenous PURA (anti-PURA$^{12D11}$) for peak calling and subsequent binding site definition. Instead, we used the PURA binding sites identified for endogenous PURA (anti-PURA$^{12D11}$) in HeLa cells to test for PURA binding in NPCs specifically at these sites. Indeed, we found that the crosslink events for endogenous PURA in NPCs enriched within the PURA binding sites identified from HeLa cells, indicating a similar RNA binding behavior in both cell types (**Figure 2C, F**).

### Supplementary Tables

#### Supplementary Table S1: Overview of the four PURA iCLIP experiments.

We performed four different PURA iCLIP experiments. In the first experiment (“PURA endo HeLa”), we used the in-house antibody anti-PURA$^{12D11}$ to map the binding sites of endogenous PURA in HeLa cells (4 biological replicates, s1-s4). In the second and third experiment (“PURA oe HeLa with anti-PURA12D11/anti-FLAG”), we overexpressed (oe) FLAG-tagged PURA in HeLa cells and immunoprecipitated PURA with two different antibodies (anti-PURA$^{12D11}$, 4 biological replicates, s1-s4, and anti-FLAG, 2 biological replicates, s1-s2). In the fourth experiment (“PURA endo NPC”), we mapped the endogenous PURA binding sites in neural progenitor cells (NPCs) using anti-PURA$^{12D11}$ (4 biological replicates, s1-s4). The table provides information on the replicates of all four experiments, including details on the experiment (replicate, cell type, antibody used and PCR cycles), the sequencing (file name, sequencing date, sequencing mode and barcode) and the obtained data (sequenced reads, uniquely mapped reads and crosslink events).

< provided as Excel file >

#### Supplementary Table S2: 4,391 PURA-bound genes.

The table lists of all 4,391 genes that harbor at least one binding site of endogenous PURA in HeLa cells. For each gene, information on the gene (gene name, ENSEMBL gene ID, gene type; GENCODE release 31, genome version GRCh38.p12) are given. Additionally, the genomic coordinates of the strongest PURA binding site (chromosome, start, end, strand) are given together with the PureCLIP score of the nucleotide in the center of the binding site (PureCLIP score of center), the number of crosslink events in the binding site center (number of crosslinks in center), the number of crosslink events in the complete binding site for the 4 replicates of the endogenous PURA iCLIP in HeLa cells (crosslinks - s1-4) and the transcript region that the binding site falls into (BS region). BS, binding site.

< provided as Excel file >

#### Supplementary Table S3: Enriched REACTOME and GO cellular component terms for 4,391 PURA-bound genes.

The tables list all significantly enriched REACTOME (sheet “REACTOME”) or Gene Ontology (GO) cellular compartment terms (sheet “GO – cellular components") for the 4,391 PURA-bound genes from functional enrichment analysis using the hypeR package (version 1.9.1, “hypergeometric” mode; FDR < 0.1). All genes with at least one PURA crosslink event served as background. Given are the test statistics from hypeR (i.e., pval; fdr; “signature”, number of target genes found in any term; “geneset”, number of all genes belonging to term; “overlap”, number of signature genes in gene set, “background”, number of background genes), the gene names of all hits per term (“hits") and the ratio of hits per term (“overlap”/”geneset”).

< provided as Excel file >

#### Supplementary Table S4: Differential RNA expression upon *PURA* knockdown.

The table lists all 3,415 genes with significant changes in RNA levels in *PURA* knockdown (KD) vs. control HeLa cells (false discovery rate [FDR] < 0.01). For each gene, ENSEMBL gene ID and HGNC symbol are given together with the test statistics from differential expression analysis with DESeq2 (baseMean, mean of normalized read counts across replicates; log2FoldChange, *PURA* KD over control; lfcSE, stat, test statistics; pvalue, padj, adjusted *P* value, FDR). The table further specifies whether the gene harbors at least one binding site of endogenous PURA in HeLa cells (PURA binding) and if so, provides the genome coordinates of the strongest PURA binding site within this gene (BS_seqnames*, start*, end*, width*, strand*) as well as the PureCLIP score of the binding site center (score*) and the transcript region of the binding site (BS_region*). HGNC, HUGO Gene Nomenclature Committee; BS, binding site.

< provided as Excel file >

#### Supplementary Table S5: Differential protein abundance upon *PURA* knockdown.

The table lists all 995 proteins with significant changes in protein abundance that were identified in shot-gun proteomics measurements of *PURA* knockdown and control HeLa cells (false discovery rate [FDR] < 0.05). The table summarizes the following information. For each protein, ENSEMBL gene ID and gene name are given together with the test statistics from differential expression analysis with limma (logFC, *PURA* KD over control; AveExpr, average expression; t, test statistics; P.Value; adj.P.Val, adjusted *P* value) followed by *P* value scaling by peptide counts with DEqMS (count, number of unique peptides found; sca.t; sca.P.Value; sca.adj.pval). In addition, the table lists the RNA expression changes in *PURA* KD vs. control of the corresponding RNAs. The test statistics from differential expression analysis with DESeq2 (baseMean, mean of normalized read counts across replicates; log2FoldChange, *PURA* KD over control; lfcSE, stat, test statistics; pvalue, padj, adjusted *P* value, FDR) are given. The table further specifies whether the gene harbors at least one binding site of endogenous PURA in HeLa cells (PURA binding) and if so, provides the genome coordinates of the strongest PURA binding site within this gene (BS_seqnames*, start*, end*, width*, strand*) as well as the PureCLIP score of the binding site center (score*) and the transcript region of the binding site (BS_region*).

< provided as Excel file >

### Supplementary Table S6: Enriched REACTOME and GO cellular component terms for 3,415 differentially expressed RNAs in *PURA* knockdown.

The tables list all significantly enriched REACTOME (sheet "REACTOME") or Gene Ontology (GO) cellular compartment terms (sheet "GO – cellular components") for the genes of the 3,415 differentially expressed RNAs in *PURA* knockdown. Functional enrichment analysis is performed in R using the "hypergeometric" mode (FDR < 0.1) of the hypeR package (version 1.9.1) with all genes with at least one PURA crosslink event as background. Given are the test statistics from hypeR (pval; fdr; "signature", number of target genes found in any term; "geneset", number of all genes belonging to term; "overlap", number of signature genes in geneset, "background", number of background genes), the gene names of all hits per term ("hits") and the ratio of hits per term ("overlap"/"geneset"). Enrichment analysis was performed once for RNAs downregulated in *PURA* knockdown (n = 1752, FDR < 0.01, log2foldchange < 0) and once for RNAs upregulated in *PURA* knockdown (n = 1663, FDR < 0.01, log2foldchange > 0), marked in the column "Regulation in PURA KD/CTRL" as "down" and "up", respectively.

< provided as Excel file >

### Supplementary Table S7: Enriched REACTOME and GO cellular component terms from 995 differentially expressed proteins in *PURA* knockdown.

The tables list all significantly enriched REACTOME (sheet "REACTOME") or Gene Ontology (GO) cellular compartment terms (sheet "GO – cellular components") for the genes of the 995 differentially expressed proteins in *PURA* knockdown. Functional enrichment analysis is performed in R using using the "hypergeometric" mode (FDR < 0.1) of the hypeR package (version 1.9.1) with all genes with at least one PURA crosslinked nucleotide as background. Given are the test statistics from hypeR (pval; fdr; "signature", number of target genes found in any term; "geneset", number of all genes belonging to term; "overlap", number of signature genes in geneset, "background", number of background genes), the gene names of all hits per term ("hits") and the ratio of hits per term ("overlap"/"geneset"). Enrichment analysis was performed once for proteins downregulated in *PURA* knockdown (n = 406, FDR < 0.01, log2foldchange < 0) and once for proteins upregulated in *PURA* knockdown (n = 589, FDR < 0.01, log2foldchange > 0), marked in the column "Regulation in PURA KD/CTRL" as "down" and "up", respectively.

< provided as Excel file >

### Supplementary Table S8: List of qPCR primers used.

| Primer name | Sequence | Target gene |
|---|---|---|
| PURA_qPCR_F_ mouse_temp | GCCAAGCTCATCGACGACTACG | *PURA* |
| PURA_qPCR_R_ mouse_temp | CTCACTCGCATAAACACGCCGT | *PURA* |
| PURA_ALA_2_F | CCTTACTCTCTCCATGTCAGTG | *PURA* |
| PURA_ALA_2_R | CAATGGTCTGGCCCTGC | *PURA* |
| RPL32-F- getprime | AAATTAAGCGTAACTGG CGG | *RPL32* |
| PRL32-R- getprime | GTTGGGCATCAAGATCT GG | *RPL32* |
| GAPDH_ER_F | GCTCATTTCCTGGTATGACAACG | *GAPDH* |
| GAPDH_ER_R | GAGATTCAGTGTGGTGGGGG | *GAPDH* |
| OCT_4_F | CAATTTGCCAAGCTCCTGAAG | *OCT4* |
| OCT_4_R | AAAGCGGCAGATGGTCGTT | *OCT4* |
| SOX2_F | CCTCCGGGACATGATCAGCATGTA | *SOX2* |
| SOX2_R | GCAGTGTGCCGTTAATGGCCGTG | *SOX2* |
| PAX6_F | GCGGAGTTATGATACCTACACC | *PAX6* |
| PAX6_R | GAAATGAGTCCTGTTGAAGTGG | *PAX6* |
| ACSL1.F | ACTGTGCAGAAATAAGGATGTC | *ACSL1* |
| ACSL1.R | AATGGTTTCAGACCAGAATCCT | *ACSL1* |
| FOXG1.qPCR.F | GCTGGACATGGGAGATAGG | *FOXG1* |
| FOXG1.qPCR.R | GTTGATGCTGAACGAGGAC | *FOXG1* |
| MAP2.qPCR.F | GGTCACAGGGCACCTATTCA | *MAP2* |
| MAP2.qPCR.R | TGTTCACCTTTCAGGACTGC | *MAP2* |

### Supplementary Table S9: Oligonucleotides used for iCLIP experiments.

The DNA linker (L3 App) is preadenylated at the 5' end (rApp) and closed with a dideoxycytosine (ddC) to prevent concatemerization.

| Name | Sequence (5'-3') | Application |
|---|---|---|
| L3 App Linker (pre-adenylated) | /rApp/AGATCGGAAGAGC GGTTCAG/ddC/ | 3' linker |
| RT1clip2.0 primer | GGATCCTGAACCGCT | Primer for reverse transcription |
| L01clip2.0– L12clip2.0 | As described in (11) | 5' linker and barcode |
| P3_Solexa_short | CTGAACCGCTCTTCCGA TCT | Initial PCR amplification |
| P5_Solexa_short | ACACGACGCTCTTCCGA TCT | Initial PCR amplification |
| P3_Solexa | CAAGCAGAAGACGGCAT ACGAGATCGGTCTCGG CATTCCTGCTGAACCGC TCTTCCGATCT | Final PCR amplification |
| P5_Solexa | AATGATACGGCGACCAC CGAGATCTACACTCTTT CCCTACACGACGCTCT TCCGATCT | Final PCR amplification |

### Supplementary Table S10: Oligonucleotides used for EMSA experiments.

| Target name | Sequence (5´-3´) | Length |
|---|---|---|
| *CLTA* | AATGCCAGGGAGAACACAGTTGAAGGAAGGAAACATGCAATCACAAACAACTATAGTGAGTCGTATTA | 68 |
| *CLTC* | CTAAAACACCACAATATCCAACATACACAAACCTCAGGGAAGGGTTAGTAAACACACACAAGACTATAGTGAGTCGTATTA | 81 |
| *COX7C* | agcaaatgcagatccaaagtacaaacacatcttagctagtaacgaccacttgCTATAGTGAGTCGTATTA | 70 |
| *EEF2* | GGGGCAAAAGCCACTGCGGGCCATGTACCCAAATAAACCTCTTAATGCGTTTCTATAGTGAGTCGTATTA | 70 |
| *LSM14A* | tccaaatgtttcagcatttaatcttctttcttcagcccaggttggtctccgttctctaCTATAGTGAGTCGTATTA | 76 |
| *NDUFS5* | TCCAGTGGAGAACAGAGAACAGGAAAATCAGCCTCCAGACATCAGCAGCTCTATAGTGAGTCGTATTA | 68 |
| *NEAT1* | ttacagacactgcacgccccagcaattgaggaaccagcgtggccacagcgggCTATAGTGAGTCGTATTA | 70 |
| *STARD7* | acacacaTATTCTCTCTCTCTCTTATGCACACATCCATCCACATCCCAACAATTCTATAGTGAGTCGTATTA | 72 |
| MF0677 51mer | GCATTTTAAGTAAACTTTTCTCTCCCTCCACCACCTCCAAAAGAGAAAACACTATAGTGAGTCGTATTA | 69 |
| MF0677 24mer | CTTTTCTCTCCCTCCACCACCTCCCTATAGTGAGTCGTATTA | 42 |
| *ASH1 E3* 51mer | ATTGTTTCGTGATAATGTCTCTTATTAGTTGAAAGAGATTCAGTTATCCATCCTATAGTGAGTCGTATTA | 70 |

#### Supplementary Table S11: List of primary antibodies used.

For each antibody, the clonality and source animal are listed together with the dilution for the respective applications (WB, Western blot; IF, immunofluorescence; IP, immunoprecipitation).

| Antibody | Clonality/source | Dilution/application | Source |
|---|---|---|---|
| Anti-H3 (ab1791) | Polyclonal / rabbit | 1:10000 / WB | Abcam |
| Anti-PURA$^{12D11}$ | Monoclonal / rat | 1:10 / WB<br>Undiluted / IF<br>Undiluted / IP | HMGU antibody core facility |
| Anti-Tubulin clone DM1A (2999783) | Monoclonal / mouse | 1:1000 / WB | Sigma Aldrich |
| Anti-DCP1A (# PA5-82478) | Polyclonal / rabbit | 1:1000 / IF<br>1:500 / WB | Thermo Fisher Scientific |
| Anti-LSM14A (HPA017961) | Polyclonal / rabbit | 1:1000 / IF<br>1:500 / WB | Sigma Aldrich |
| Anti-IL6ST (MAB228) | Monoclonal / mouse | 1:250 / IF | R&D systems |
| Anti-p62 lck ligand (#610832) | Monoclonal / mouse | 1:1000 / WB | BD Biosciences |
| Anti-GAPDH hFab Rhodamine | Monoclonal / HuCAL | 1:5000 / WB | BioRad |
| Anti-PARP1 | Polyclonal / rabbit | 1:5000 / WB | Sigma Aldrich |
| Anti-GST | Monoclonal / rat | 1:10 / WB | HMGU antibody core facility |
| Anti-DDX6 (HPA024201) | Polyclonal / rabbit | 1:1000 / WB | Sigma Aldrich |
| Anti-IL6ST (sc-376280) | Monoclonal / mouse | 1:250 / WB | Santa Cruz Biotechnology |

### Supplementary Table S12: List of secondary antibodies used.

For each antibody, the clonality and source animal are listed together with the dilution for the respective applications (WB, Western blot; IF, immunofluorescence).

| Antibody | Clonality/source | Dilution/application | Source |
|---|---|---|---|
| Mouse-anti-rat IgG2a-HRP | Polyclonal / mouse | 1:1000 / WB | HMGU antibody core facility |
| Donkey-anti-rat AlexaFluor 488 (A21208) | Polyclonal / donkey | 1:1000 / IF | Thermo Fisher Scientific |
| Donkey-anti-rat AlexaFluor 555 (ab150154) | Polyclonal / donkey | 1:1000 / IF | Abcam |
| Goat-anti-rabbit IgG H&L (HRP) (ab6721) | Polyclonal / goat | 1:100.000 / WB | Abcam |
| Goat-anti-mouse HRP (G-21040) | Polyclonal / goat | 1:20.000 / WB | Thermo Fisher Scientific |
| Goat anti-mouse IgG - Starbright Blue 520 | Polyclonal / goat | 1:5000 / WB | BioRad |
| Goat-anti-rat-647 (ab150167) | Polyclonal / rat | 1:1000 / IF | Abcam |
| Donkey-anti-rabbit-555 (A31572) | Polyclonal / rabbit | 1:1000 / IF | Thermo Fisher Scientific |
| Goat-anti-mouse-488 (A32723) | Polyclonal / goat | 1:1000 / IF | Invitrogen |
